# Dynamic assembly of malate dehydrogenase-citrate synthase multienzyme complex in the mitochondria

**DOI:** 10.1101/2025.06.16.659985

**Authors:** Joy Omini, Inga Krassovskaya, Taiwo Dele-Osibanjo, Connor Pedersen, Toshihiro Obata

## Abstract

The tricarboxylic acid (TCA) cycle enzymes malate dehydrogenase (MDH1) and citrate synthase (CIT1) form a multienzyme complex, referred to as a metabolon, that channels intermediate oxaloacetate between their reaction centers. Given that the MDH1-CIT1 metabolon enhances pathway reactions in vitro, its dynamic assembly is hypothesized to contribute to TCA cycle regulation in response to cellular metabolic demands. Here, we demonstrated that yeast mitochondrial MDH1 and CIT1 dissociated when aerobic respiration was suppressed by the Crabtree effect and associated when the respiratory activity was enhanced by acetate.

Pharmacological TCA cycle inhibition dissociated the complex, whereas electron transport chain inhibition enhanced the interaction. The multienzyme complex assembly was related to the mitochondrial matrix acidification and oxidation, as well as cellular levels of malate, fumarate, and citrate. These factors significantly affected the MDH1-CIT1 complex affinity in vitro. Especially, variations in buffer pH within the physiological pH range between 6.0 and 7.0 in the mitochondrial matrix significantly impacted the MDH1-CIT1 affinity. These results demonstrate the dynamic association and dissociation of the MDH1-CIT1 metabolon and its relationship with respiratory activity, supporting metabolon dynamics as an integral factor in metabolic regulation governed by multiple factors such as mitochondrial pH and metabolite levels.

## Introduction

Enzymes catalyzing sequential reactions can interact to form a multienzyme complex, often called a ‘metabolon,’ which channels the metabolic intermediate within the complex. Metabolite channeling can mediate pathway reactions by concentrating the reaction intermediates near the enzyme active site and sequestering them from competing reactions (Obata, 2020; Spivey and Ovádi, 1999; Wheeldon et al., 2016). Thus, the dynamic assembly of multienzyme complex is believed to quickly regulate cellular metabolic flux by changing their degree of association and dissociation without involving time-consuming and resource-demanding protein synthesis, degradation, and modification (Halper and Srere, 1977; Morgunov and Srere, 1998; Obata, 2019; Omini et al., 2024; Srere, 1985; Sumegi et al., 1993). However, limited experimental evidence shows metabolon dynamics and the mechanisms regulating its association.

The tricarboxylic acid (TCA) cycle multienzyme complex composed of malate dehydrogenase (MDH) and citrate synthase (CS) is conserved in all organisms analyzed so far, including animals, bacteria, yeast, and plants (Meyer et al., 2011; Morgunov and Srere, 1998; Robinson et al., 1987; Zhang et al., 2017). MDH and CS catalyze the key steps in respiratory metabolism as their reactions comprise the carbon entry steps from the glycolytic processes as malate and acetyl-CoA, respectively (Sweetlove et al., 2010). MDH-CS complex is considered a dynamic protein complex (Bulutoglu et al., 2016; Omini et al., 2021) that protects the channeled intermediate, oxaloacetate, from degradation and competing pathways in the bulk phase of the cell (Bulutoglu et al., 2016).

Importantly, MDH-CS complex formation and oxaloacetate channeling are considered essential for the forward TCA cycle flux to occur since MDH forward reaction to synthesize oxaloacetate is thermodynamically unfavorable in physiological conditions (Fukuda et al., 2008). These findings suggest that the dynamic association and dissociation of the MDH-CS multienzyme complex play a role in coordinating the TCA cycle and associated metabolic pathways. To address this hypothesis, this study focuses on demonstrating the in vivo dynamics of the MDH-CS complex in relation to respiratory states, a crucial aspect of achieving metabolic regulation. Another key requirement, the direct impacts of the metabolon on pathway flux, will be addressed in a separate study.

This study utilized the yeast mitochondrial MDH (MDH1)-CS (CIT1) multienzyme complex as the model. Budding yeast, *Saccharomyces cerevisiae*, dramatically rearranges its central carbon metabolism in response to changes in nutrient availability and stress conditions (Causton et al., 2001; Rep et al., 1999; Rolland et al., 2002). Remarkably, yeast respiratory metabolism, involving aerobic respiration and fermentation, is highly adaptable to substrate availability (Pfeiffer and Morley, 2014). When oxygen and non-fermentable respiratory substrates, such as acetate, are abundant, aerobic respiration is upregulated, with increased carbon flux through the TCA cycle (Gerstmeir et al., 2003; Lee et al., 2011). In the presence of fermentable sugars, such as glucose and fructose, the fermentation pathway is upregulated, and aerobic respiration is repressed, even when oxygen is available (Zhang et al., 2022). Furthermore, aerobic respiration and fermentation cooperate in the presence of a poorly fermentable carbon source like raffinose (Guaragnella et al., 2013). Application of fermentable substrates to respiring yeast cells induces a substantial shift from aerobic respiration to fermentation, known as the ‘Crabtree effect,’ involving a massive transcriptional rearrangement of enzyme genes related to aerobic respiration and fermentation (Gancedo, 1998; Ronne, 1995; Thierie and Penninckx, 2010). This inducible metabolic shift in yeast makes it an ideal model for investigating the dynamic relationship between the MDH1-CIT1 multienzyme complex and TCA cycle flux. Understanding the mechanisms of the Crabtree effect is crucial for metabolic engineering applications to enhance the supply of TCA cycle intermediates for desired product synthesis (Yin et al., 2015) and for gaining insights into metabolic regulation in other eukaryotic systems, including cancer cells, which exhibit metabolic shifts similar to the Crabtree effect (Diaz-Ruiz et al., 2011).

Various allosteric regulators and environmental factors, including pH and redox state, affect MDH and CS enzyme activities (Beeckmans, 1984; Williamson and Cooper, 1980). These factors alter protein conformations, influencing the MDH-CS complex affinities. Previous in vitro studies have demonstrated that NAD^+^, malate, succinate, acetyl-CoA, α-ketoglutarate, and acidic pH enhance the MDH-CS interaction, while NADH, citrate, and basic pH weaken it (Omini et al., 2024, 2021; Tompa et al., 1987; Wu and Minteer, 2015). These allosteric regulators promote the MDH-CS interaction when the TCA cycle substrates are abundant and products are limited, indicating the role of the MDH-CS multienzyme complex in the TCA cycle feedback regulation. The electron transport chain activity affects the mitochondrial matrix pH, redox state, and ATP content and is closely related to cellular respiratory flux distributions (Santo-Domingo and Demaurex, 2012)).

Therefore, the MDH-CS multienzyme complex may associate and dissociate according to the microenvironment in the mitochondrial matrix, such as metabolite concentrations, pH, redox state, and energy levels.

We adopted a NanoBiT protein-protein interaction assay system (Dixon et al., 2016) to monitor real-time multienzyme complex interactions in living yeast cells using continuous microplate readings. The NanoBiT split-luciferase system is based on the 18 kDa NanoLUC luciferase derived from deep-sea shrimp. The small size of this reporter is designed to minimize structural and behavioral interference with the target proteins, while offering a dynamic range and brightness approximately 150-fold greater than those of conventional luciferases, enabling detection of weak, transient interactions. The NanoBiT subunits exhibit low intrinsic affinity (*K*_D_ = 190 µM) and rapid association and dissociation kinetics (*k*_on_ = 500 M^-1^s^-1^, *k*_off_ = 0.2 s^-1^), ensuring that the luminescent complex formation is dictated by the interaction of the fused proteins rather than the affinity of the tags themselves (Dixon et al., 2016). While other fluorescence-based methods, such as FLIM-FRET and BRET, are widely used for *in vivo* protein-protein interaction studies, they are limited in yeast microplate assays due to high cellular autofluorescence. The NanoBiT system overcomes these constraints, providing semi-quantitative monitoring of relative fluctuations in protein interaction levels over time across a diverse range of metabolic conditions.

In this study, the NanoBiT system examined the MDH1-CIT1 multienzyme complex association in the presence of various carbon sources and respiratory inhibitors. The results showed the relationship between respiratory activity and the dynamic MDH1-CIT1 complex assembly.

Analysis of the mitochondrial matrix microenvironment using fluorescent biosensors and cellular metabolite profiles revealed crosstalk among cellular respiratory status, the MDH1-CIT1 interaction, and the mitochondrial matrix microenvironment, including pH. These findings indicate the functional dynamics of the MDH-CS multienzyme complex and how they coordinate with the metabolic states of the TCA cycle and adjacent pathways.

## Results

### MDH1-CIT1 interaction was detected exclusively in respiratory conditions

We employed a NanoBiT split-luciferase system (Dixon et al., 2016) for semi-quantitative relative detection of the MDH1-CIT1 multienzyme complex association in living yeast cells. The codon-optimized sequences of nano-BIT subunits were integrated into the direct downstream of the MDH1 and CIT1 exons in the yeast genome by gene homologous recombination to fuse the NanoBiT subunits to the native enzymes. The NanoBiT subunits reconstitute the NanoLUC holoenzyme to produce luciferase signals when MDH1 and CIT1 are in proximity. The affinity between the NanoBiT subunits is very low and does not stabilize the interactions of the tagged proteins (Dixon et al., 2016). The integration of NanoBiT subunits had no significant effect on the cellular growth rate or cellular MDH and CS enzyme activities (Figure 1 – figure supplement 1). Although the MDH1-CIT1 reporter line did not show a detectable luciferase signal when it was grown in a fermentation condition (glucose-containing SD media; SD-Gluc), it showed a substantial luciferase signal in mixed-respiration (raffinose-containing SD media; SD-Raff) and respiration (acetate-containing SD media; SD-Acet) conditions (Figure 1A). These results indicate that MDH1 and CIT1 interact under respiratory conditions. The MDH1-CIT1 interaction can be related to the oxidative respiration and the TCA cycle activity since oxygen consumption was detected only in SD-Raff and SD-Acet media but not in the SD-Gluc medium (Figure 1B), as previously reported (Klein et al., 1998; Polakis and Bartley, 1965; Rep et al., 1999; Rolland et al., 2002; Strittmatter, 1957). However, MDH1 and CIT1 expressions were downregulated in SD-Gluc growing cells near the detection limit (Figure 1C), making it difficult to conclude if the undetectable MDH1-CIT1 interaction was due to complex dissociation or low protein levels.

**Figure 1.**
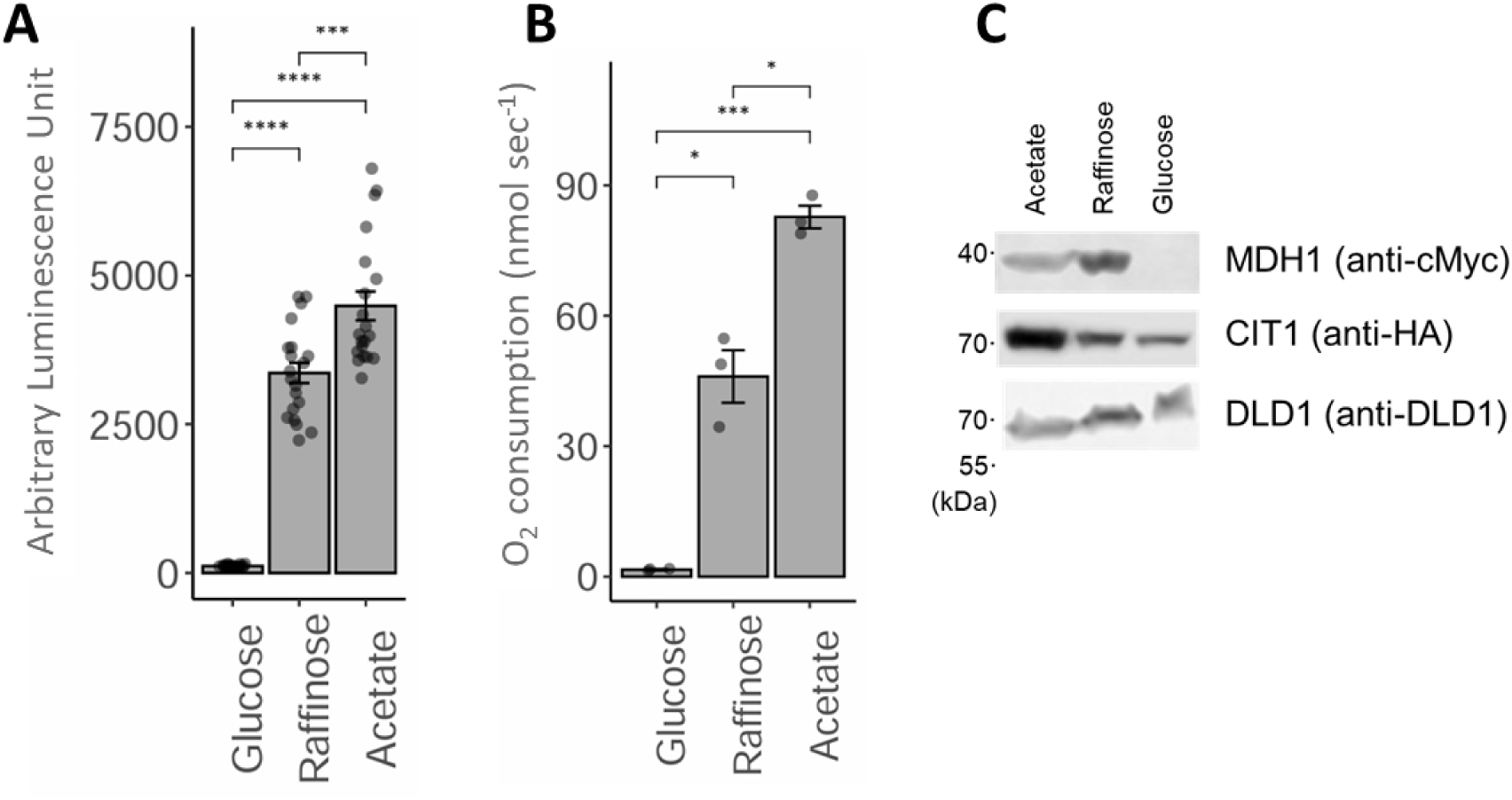
MDH1-CIT1 interaction under respiration, fermentation, and mixed-respiration conditions. Yeast cells were grown in the minimum media containing acetate (SD-Acet), glucose (SD-Gluc), and raffinose (SD-Raff) to the exponential growth phase. **(A)** Luciferase signal indicating MDH1-CIT1 complex interaction (N=20). **(B)** Cellular oxygen consumption rate (N=3). **(C)** MDH1 and CIT1 protein levels detected by Western blotting. Numbers on the left indicate the position of the molecular weight markers. DLD1 is a loading control of mitochondrial protein. Full gel images are in Figure 1 – figure supplement 2. In A and B, data are presented as mean ± SEM. and the differences between conditions were tested by Student’s *t*-test. Asterisks indicate significant differences (*, *p*<0.05; **, *p*<0.01; ***, *p*<0.001; ****, *p*<0.0001; ns, not significant).

## Crabtree induction reduced the MDH1-CIT1 interaction

To test the relationship between respiratory activity and the MDH1-CIT1 complex association when MDH1 and CIT1 enzymes are abundant in the cell, we monitored the time course of the MDH1-CIT1 interaction after a rapid shift from aerobic to anaerobic respiration via the Crabtree effect. The Crabtree effect was induced by applying 2% glucose to the SD-Raff-grown MDH1-CIT1 NanoBiT reporter line (Figure 2, Figure 2 – figure supplement 1). The luciferase signals only slightly changed for 30 min upon glucose application, followed by a steep decline. In contrast, the control cells retained the initial signal level for 100 min (Figure 2A, Figure 2 – figure supplement 1D). Decrease in the MDH1-CIT1 interaction was validated by a co-immunoprecipitation assay at 1.5 and 2.5 hours after the glucose application (Figure 2 – figure supplement 1A). Other Crabtree inducers, such as fructose and sucrose, also reduced the MDH1-CIT1 interaction, while the addition of non-fermentable sugars, including galactose, caused no significant change in MDH1-CIT1 interaction (Figure 2 – figure supplement 2A-D, Figure 2 – figure supplement 3), likely due to raffinose in the media suppressing its effects. The signal decline following glucose application was partially reversed by co-application with a fermentation inhibitor, 100 mM phosphate (van Urk et al., 1989; Figure 2A). Phosphate co-application slowed the decrease of luciferase signal compared to the glucose-applied cells, and the signal was not statistically significantly lower than the control until 80 min after application. The oxygen consumption rate significantly decreased when glucose was added to SD-Raff-grown cells but was slightly recovered by co-application of glucose with phosphate (Figure 2 – figure supplement 2E). These results indicate that the respiratory suppression by the Crabtree effect is related to MDH1-CIT1 complex disruption.

**Figure 2.**
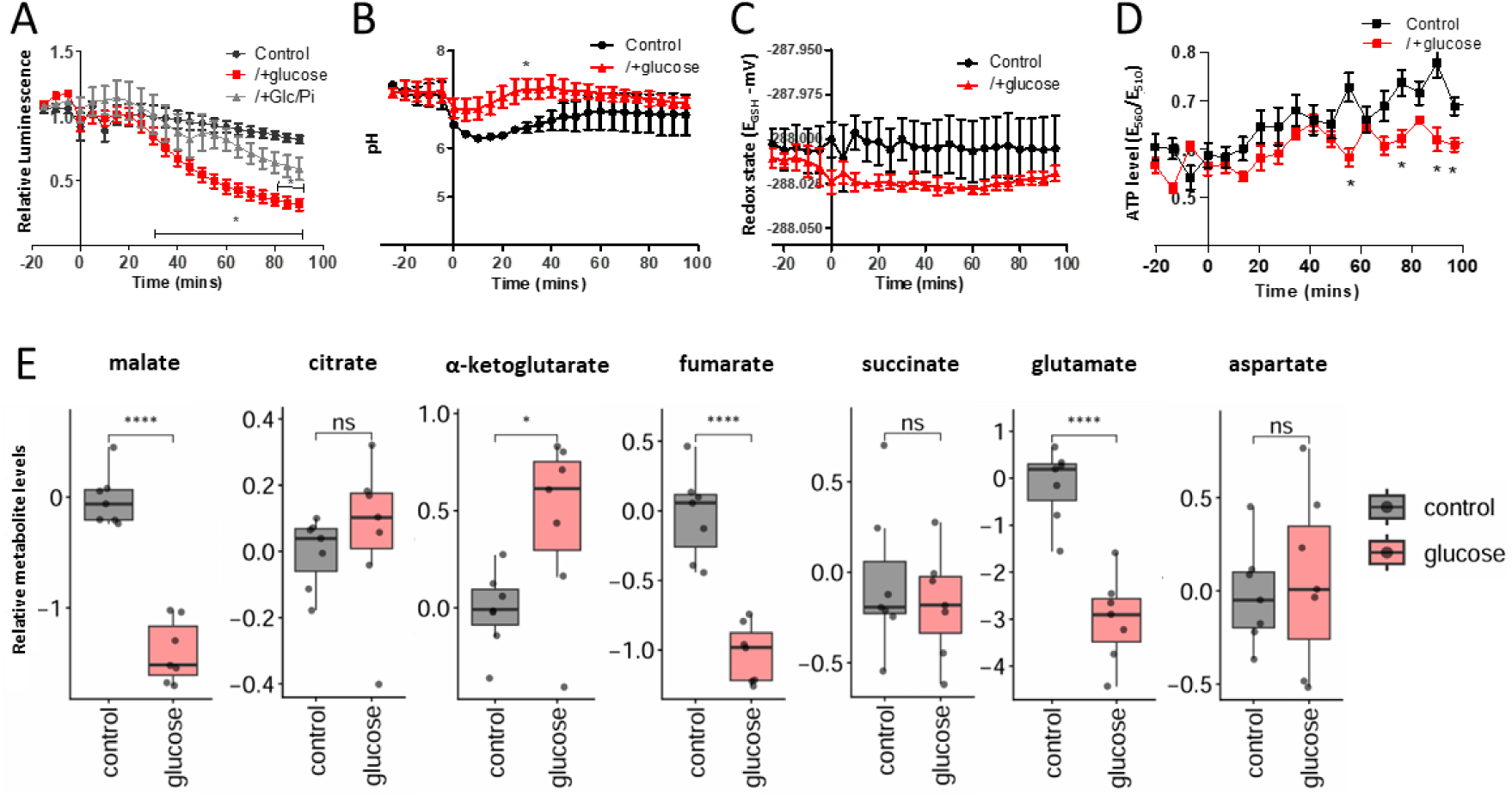
MDH1-CIT1 complex association, mitochondrial microenvironments, and cellular metabolite levels during Crabtree effect induction. Cells were cultured in fresh SD-Raff media in the control condition (black). The Crabtree effect was induced by the 2% glucose application to the SD-Raff-grown cells at 0 min (red). **(A)** NanoBIT signal indicating MDH1-CIT1 interaction. Relative luciferase unit (RLU) was calculated by normalizing the luciferase signals tothe average signals during three pre-treatment time points. SD-Raff-grown cells were also co-treated with 2% glucose and a fermentation inhibitor, 100 mM phosphate, at 0 min (gray). **(B)** Mitochondrial matrix pH. **(C)** Mitochondrial matrix redox states as GSH/GSSG equivalent (mV). **(D)** Mitochondrial matrix ATP level indicated by the ratio between 560 and 510 nm emission signals. All data in A-D are presented as mean ± s.d. (N=4). **(E)** Cellular metabolite levels at 80 min. The boxes, lines, error bars, and points indicate interquartile range, median, minimum, and maximum values, and outliers, respectively (N=7). Statistical differences against the control samples were assessed using the Student’s *t*-test at each time point. Asterisks indicate significant differences (*, *p*<0.05; **, *p*<0.01; ***, *p*<0.001; ****, *p*<0.0001; ns, not significant).

MDH1 protein level monitored with the yeast strain expressing MDH1 fused with full-length NanoLUC luciferase showed a 20% decline within 100 min upon glucose addition (Figure 2, Figure 2 – figure supplement 1B). Western blotting also showed a slight decrease in the MDH1 and CIT1 levels (Figure 2 – figure supplement 1C). The time course of the MDH1-CIT1 NanoBiT signal was inconsistent with that of MDH1-NanoLUC, and the 20% decline in protein content cannot fully explain the decrease of MDH1-CIT1 signal by over 50% within 100 min after glucose addition (Figure 2A, Figure 2 – figure supplement 1B). The reduction of MDH1-CIT1 interaction was not accompanied by the MDH1 and CIT1 abundance in a repeated experiment (Figure 2 – figure supplement 1D-E). The interaction index, calculated by normalizing the NanoBiT signal to MDH1 and CIT1 abundances measured with MDH1-NanoLUC and CIT1-NanoLUC (see Materials and Methods), showed reduced interaction after 60 min of glucose application.

Therefore, the decline in the MDH1-CIT1 NanoBiT signal reflects the MDH1-CIT1 complex dissociation co-occurring with the Crabtree effect, even though we cannot exclude a partial effect of protein degradation (Litsios et al., 2018).

Oxidative respiration significantly influences the microenvironments in the mitochondrial matrix (Beeckmans, 1984; Williamson and Cooper, 1980). To assess if the MDH1-CIT1 interaction decline is related to the changes in the microenvironments of the mitochondrial matrix by the Crabtree effect, we monitored the mitochondrial matrix pH, ATP concentration, and redox state by expressing fluorescent biomarkers, pHluorin, mito-GoAteam2, and mito-roGFP1, respectively, in the mitochondrial matrix of the MDH1-CIT1 reporter line (Figure 2 – figure supplement 4A-D).

Mitochondrial matrix pH in the SD-Raff-grown cells was 7.2 and temporally declined to 6.8 in the first 25 min of glucose application. The control cell matrix pH gradually decreased to 6.2 and remained lower than in glucose-applied cells (Figure 2B). The alterations of the MDH1-CIT1 interaction and mitochondrial matrix parameters in the control condition are not due to carbon starvation, since 2% raffinose application had no significant effect (Figure 2 – figure supplement 4E). The redox state stayed around -288.0 mV in both the control and glucose cells (Figure 2C). The ATP level slightly increased during the experimental period of 100 min in both the control and glucose cells, with slightly but significantly lower levels in the glucose than in the control cells from 55 min to 100 min (Figure 2D). Thus, mitochondrial matrix pH and ATP levels were maintained after glucose was applied to cells, while they decreased and increased in control cells during the experimental period, indicating that mitochondrial pH and ATP levels are associated with MDH-CS complex dissociation during the Crabtree induction.

Our previous in vitro study showed that metabolite abundance influences the MDH-CS complex interaction (Omini et al., 2021). To assess the possible roles of metabolite accumulation in controlling MDH1-CIT1 complex interaction, we determined the cellular levels of 38 metabolites (Data S1) by gas chromatography-mass spectrometry (GC-MS). We focus here on the intermediates of the TCA cycle and related pathways (Figure 2E). At 80 min after 2% glucose application to SD-Raff-grown cells, malate, citrate, fumarate, and glutamate levels decreased, while α-ketoglutarate content increased significantly compared to the control cells (Figure 2G). Thus, the glucose-induced shift from aerobic respiration to fermentation altered the intracellular metabolite profile in SD-Raff-grown cells, potentially influencing the MDH1-CIT1 affinity.

## MDH1-CIT1 complex interacts in relation to the TCA cycle activity

We further assessed the relationship between the MDH1-CIT1 complex association and TCA cycle activity. Acetate supplies acetyl-CoA to the TCA cycle and enhances its activity (Cavero et al., 2003). The application of 1% sodium acetate gradually increased the MDH1-CIT1 interaction while MDH1 and CIT1 protein abundance slightly reduced (Figure 3A, Figure 3 – figure supplement 1 A-D). The TCA cycle activation by ethanol (Schüller, 2003) also enhanced the complex association (Figure 3 – figure supplement 1E). However, it is accompanied by an increase in MDH1 and CIT1 abundance (Figure 3 – figure supplement 1F&G), making the effects of ethanol on the complex association unclear (Figure 3 – figure supplement 1H). On the other hand, TCA cycle inhibition by 5 mM sodium arsenite, which impedes α-ketoglutarate dehydrogenase (Lee et al., 2011), led to a decline in MDH1-CIT1 interaction within 10 min (Figure 3B, Figure 3 – figure supplement 1 I-L). Aminooxyacetate (AOA) is an aminotransferase inhibitor and reduces TCA cycle activity by blocking the malate-aspartate NADH shuttle, which intersects with the cycle (Borst, 2020; Eto et al., 1999; Molinié et al., 2022). The MDH1-CIT1 interaction gradually reduced following 0.5 mM AOA application (Figure 3C, Figure 3 – figure supplement 1M-P). These results support that the MDH1-CIT1 multienzyme complex association is related to TCA cycle activity.

**Figure 3.**
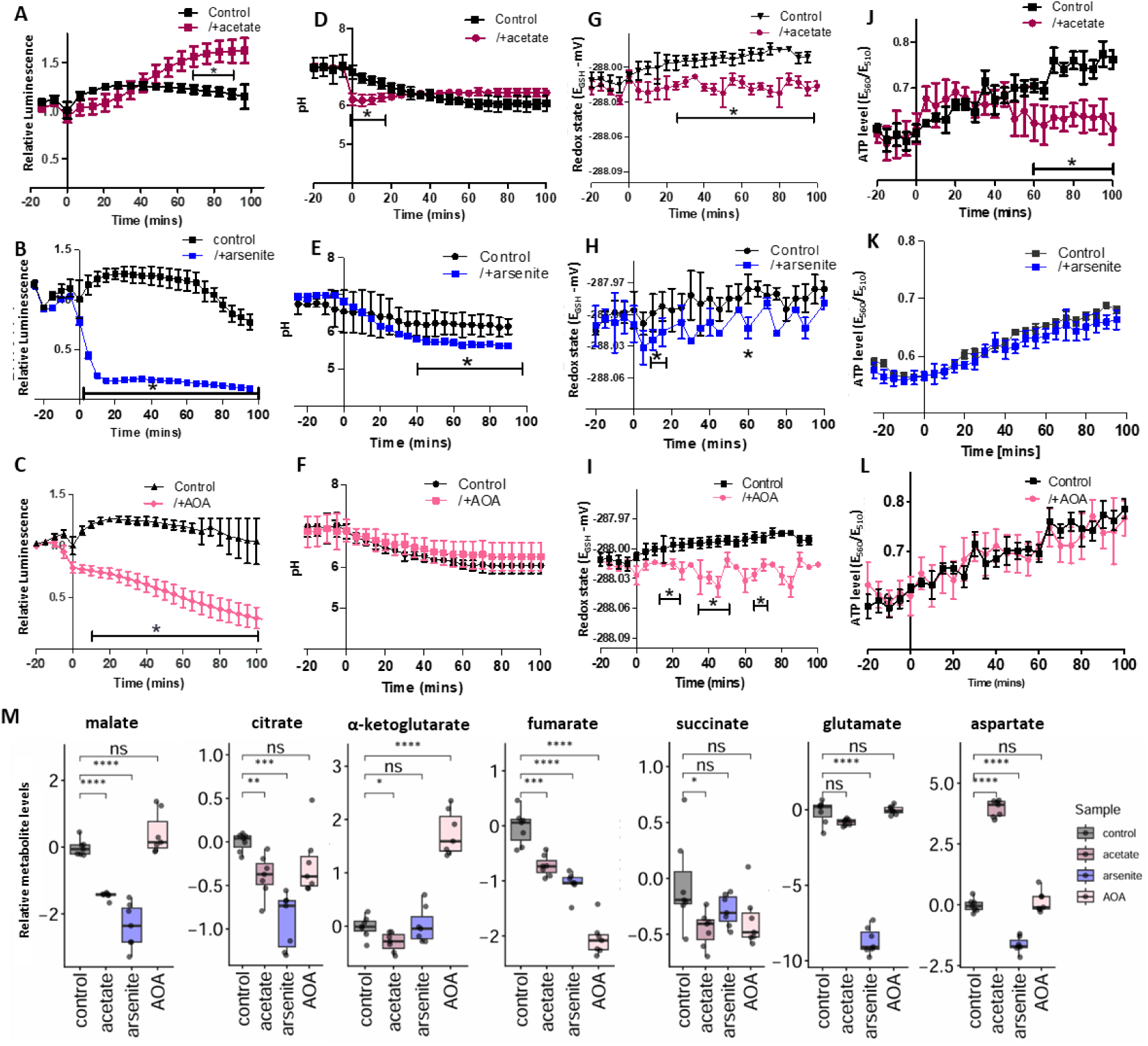
MDH1-CIT1 complex association, mitochondrial matrix microenvironments, and cellular metabolite levels following TCA cycle activation and inhibition. Cells were cultured in SD-Raff media in the control condition (black). The TCA cycle activator (acetate, dark red) and inhibitors (arsenite, blue, and aminooxyacetate (AOA), pink) were applied at 0 min. **(A-C)** NanoBiT signal indicating MDH1-CIT1 interaction. Relative luciferase unit (RLU) was calculated by normalizing the luciferase signals by the average signals during three pre-treatment time points. **(D-F)** Mitochondrial matrix pH in control cells (black) and cells treated with acetate (dark red), arsenite (blue) and AOA (pink). **(G-I)** Mitochondrial matrix redox states as GSH/GSSG equivalent (mV). **(D, H, L)** Mitochondrial matrix ATP level indicated by the ratio between 560 and 510 nm emission signals of mito-GoATeam2 sensor. All data in A-L are presented as mean ± s.d. (N=4). **(M)** Cellular metabolite levels after 80 min of treatment. The boxes, lines, error bars, and points indicate interquartile range, median, minimum, and maximum values, and outliers, respectively (N=7). Statistical differences against the control samples were assessed using the Student’s *t*-test at each time point. Asterisks indicate significant differences (*, *p*<0.05; **, *p*<0.01; ***, *p*<0.001; ****, *p*<0.0001; ns, not significant).

Mitochondrial matrix pH declined to 6.1 immediately after acetate application (Figure 3D). Mitochondrial pH went slightly lower than that of the control cells after 40 min of TCA cycle inhibition by arsenite (Figure 3E), while AOA application had no effect on mitochondrial pH (Figure 3F). The mitochondrial matrix redox state was gradually increased in the control cells during the experimental period, while it was kept constant in the cells treated by acetate, arsenite, and AOA (Figure 3G-I), resulting in the significantly lower mitochondrial matrix redox state in the treated cells in the latter half of the experimental period. Mitochondrial ATP level was decreased by acetate treatment after 40 min (Figure 3J), although arsenite and AOA showed no effect on the mitochondrial ATP levels within the experimental period (Figure 3J&K). These changes in mitochondrial microenvironments did not show clear, direct relationships with the MDH1-CIT1 interaction.

We evaluated the cellular metabolite profile after treatment of SD-Raff grown cells for 80 min with 1% acetate, 5mM arsenite, and 0.5 mM AOA to determine the relationship between change in metabolite levels and the MDH1-CIT1 complex interaction (Figure 3M). Acetate application significantly reduced malate, α-ketoglutarate, fumarate, succinate, and glutamate levels, while aspartate abundance significantly increased. Inhibition of the TCA cycle by arsenite significantly reduced the levels of all TCA cycle metabolites other than α-ketoglutarate. Aspartate and glutamate were also decreased, and glutamate was almost depleted in the cell (Figure 3M). AOA significantly decreased citrate, fumarate, and succinate levels while it increased α-ketoglutarate abundance (Figure 3M). Reduction of TCA cycle organic acids by the TCA cycle inhibition may be related to the reduction in the MDH1-CIT1 interaction.

## Electron transport chain inhibition enhanced the MDH1-CIT1 interaction, coinciding with matrix acidification

Mitochondrial electron transport chain (ETC) activity is directly related to the mitochondrial microenvironments (Selivanov et al., 2008). To further assess the effects of mitochondrial matrix microenvironments on MDH1-CIT1 complex interaction, we treated SD-Raff-grown cells with ETC inhibitors. Inhibition of ETC complex II, III, IV, and V with 20 mM malonate, 10 µM antimycin, 0.5 mM cyanide, and 1 mM oligomycin, respectively, significantly reduced oxygen consumption of SD-Raff-grown cells (Figure 4 – figure supplement 1), showing their effects on respiration. MDH1-CIT1 interaction was enhanced by ETC inhibitors with minor alterations of MDH1 and CIT1 protein levels (Figure 4A-C, Figure 4 – figure supplement 2), whereas the complex V (ATP synthase) inhibitor exerted no effect on MDH1-CIT1 interaction and mitochondrial matrix microenvironment in this experimental condition (Figure 4 – figure supplement 3A-D). Complex II inhibition slightly and temporally enhanced the MDH1-CIT1 interaction, although the increase was not statistically significant compared to the control (Figure 4A, Figure 4 – figure supplement 1B-E). The MDH1-CIT1 interaction was enhanced in the first 20 min of complex IV inhibition and reduced to the basal level afterward (Figure 4B, Figure 4 – figure supplement 2E-H). Complex III inhibition by antimycin slowly increased MDH1-CIT1 interaction, and the signal increase peaked at 60 min of treatment (Figure 4C; Figure 4 – figure supplement 2I-L).

**Figure 4.**
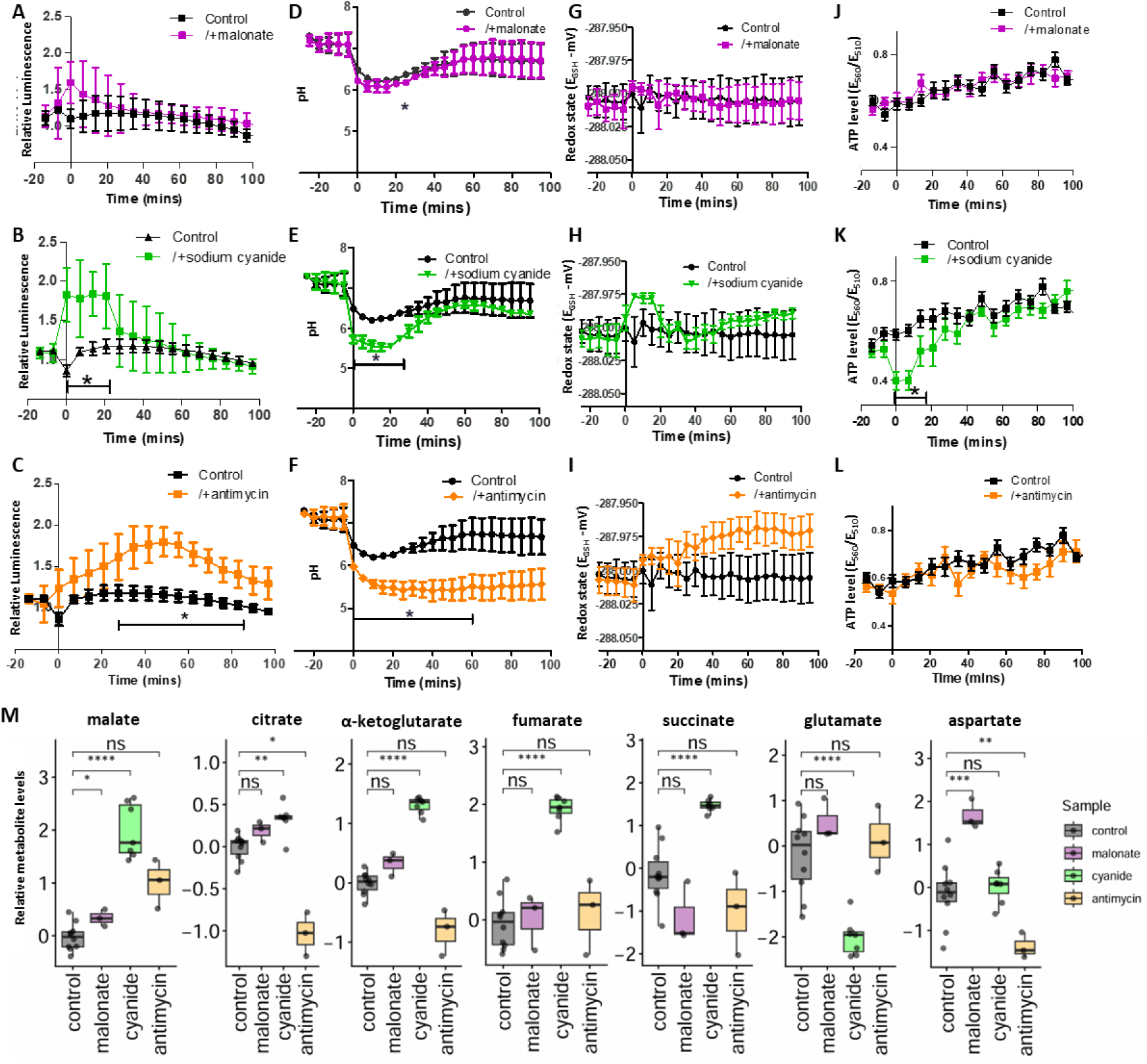
MDH1-CIT1 complex association, mitochondria microenvironments, and cellular metabolite levels following mitochondrial electron transport chain (ETC) inhibition. Cells were cultured in SD-Raff media in the control condition (black). The ETC inhibitors for complex II (malonate, purple, **A-D**), complex IV (cyanide, green, **E-H**), and complex III (antimycin, orange, **I-L**) were applied at 0 min. **(A, E, I)** NanoBiT signal indicating MDH1-CIT1 interaction. Relative luciferase unit (RLU) was calculated by normalizing the luciferase signals by the average signals during three pre-treatment time points. **(B, F, J)** Mitochondrial matrix pH. **(C, G, K)** Mitochondrial matrix redox states as GSH/GSSG equivalent (mV). **(D, H, L)** Mitochondrial matrix ATP level indicated by the ratio between 560 and 510 nm emission signals of mito-GoATeam2 sensor. All data in A-L are presented as mean ± s.d. **(M)** Cellular metabolite levels after 30 min for malonate and cyanide and after 80 min for antimycin treatment. The boxes, lines, error bars, and points indicate interquartile range, median, minimum, and maximum values, and outliers, respectively. Statistical differences against the control samples were assessed using the Student’s *t*-test at each time point. Asterisks indicate significant differences with *p*<0.05.

Mitochondrial matrix pH was reduced by all ETC inhibitor treatments (Figure 4D-F), although the effect was very small in malonate-treated cells (Figure 4D). The matrix pH was transiently decreased and exhibited an inverse relationship with the MDH1-CIT1 interaction (Figure 4E). The mitochondrial matrix pH was reduced to 5.5 immediately after antimycin application and remained at the same level for 100 min (Figure 4F). These treatments show minor effects on the redox state of the mitochondrial matrix (Figure 4 G-I). The matrix was gradually oxidized following antimycin treatment, but the difference relative to control cells was not statistically significant (Figure 4I). ETC inhibitions also showed relatively minor effects on mitochondrial ATP levels (Figure 4J-L), except for the reduction of ATP levels at the first 20 min of Complex IV inhibition (Figure 4K), which is accompanied by the increase of MDH1-CIT1 interaction (Figure 4B). These results indicate the relationship between MDH1-CIT1 interaction and mitochondrial matrix acidification.

Since the results indicate a relationship between mitochondrial matrix pH and the MDH1-CIT1 interaction, we have tested the effect of an uncoupler, carbonyl cyanide 3-chlorophenylhydrazone (CCCP), on this interaction (Figure 4 – figure supplement 4), which is expected to equilibrate the mitochondrial pH with the medium pH of 5.8 (Rosado et al., 2008). Treatment with 2 µM CCCP increased the MDH1-CIT1 interaction signal, suggesting that acidic pH enhances complex association (Figure 4 – figure supplement 4A&D). However, this result should be interpreted with caution, as CCCP treatment compromised the luciferase signal from MDH1 and CIT1 fused to full-length NanoLUC (Figure 4 – figure supplement 4B&C).

Cellular metabolite profiles were determined 30 min after malonate and cyanide treatments and 80 min after antimycin and oligomycin treatments. Complex II and V inhibition did not significantly alter cellular metabolite levels other than the malate and aspartate accumulation in malonate-treated cells (Figure 4M, Figure 4 – figure supplement 3E). Complex IV inhibition by cyanide significantly reduced citrate and glutamate levels, while malate, alpha-ketoglutarate, fumarate, and succinate levels significantly increased (Figure 4M). Complex III inhibition significantly decreased citrate and α-ketoglutarate levels and increased the malate level (Figure 4M). These results support the relationship between the levels of TCA cycle intermediates and MDH1-CIT1 interaction.

## MDH1-CIT1 interaction is affected by pH, ATP concentration, and metabolite availability in vitro

To evaluate the specific effects of microenvironments and metabolite availability on the MDH1-CIT1 interaction, we investigated the MDH1-CIT1 affinity in vitro using recombinant proteins.

Acidic pH significantly enhanced the binding affinity of the MDH1-CIT1 complex (Figure 5A) within the physiological range observed in this study (Figure 2B, Figure 3D, Figure 4D, E, F). The *K*_d_ of the MDH1-CIT1 interaction was 3.48 µM at pH 7.2, while it was one magnitude lower at acidic pH (0.223 µM at pH 6.4 and 0.033 µM at pH 6.0). We also assessed the effects of the metabolites on the MDH1-CIT1 interaction (Figure 5B, Figure 5 – figure supplement 1). We tested malate, fumarate, citrate, α-ketoglutarate, succinate, glutamate, and aspartate since their cellular levels changed when the MDH1-CIT1 interaction altered (Figure 2E, Figure 3M, Figure 4M). Citrate destroyed the MDH1-CIT1 interaction, while malate and fumarate significantly enhanced complex affinity (Figure 5B). α-ketoglutarate, succinate, and aspartate slightly enhanced MDH1-CIT1 complex affinity, although glutamate did not affect the interaction (Figure 5 – figure supplement 1A). ATP also enhanced the MDH1-CIT1 complex affinity (Figure 5 – figure supplement 1B) as observed in porcine enzymes (Omini et al., 2021). Thus, malate, fumarate, citrate, ATP levels, and pH potentially influence the MDH1-CIT1 interaction in yeast mitochondria.

**Figure 5.**
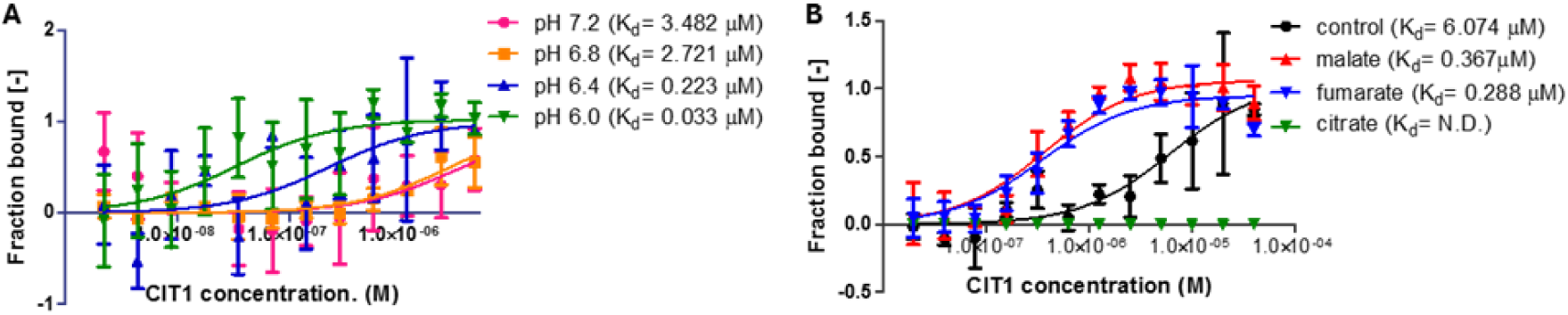
Effects of pH and metabolites on the yeast MDH1-CIT1 multienzyme complex affinity. The affinity of the MDH-CS multienzyme complex was analyzed by microscale thermophoresis (MST) using fluorescently labeled MDH1 as the target and CIT1 as the ligand. Curves represent the response (fraction bound) against CIT1 concentration. Data is presented as mean ± s.d. (N=3). **(A)** Effects of pH. The MDH1-CIT1 interaction was determined in the buffer with pH 7.2 (pink), 6.8 (orange), 6.4 (olive green), 6.0 (green), and 5.8 (blue). **(B)** Effects of 10 mM malate (red), α-ketoglutarate (green), succinate (brown), citrate (blue), aspartate (purple), glutamate (pink), and fumarate (orange). The *K*_d_ values of MDH1-CIT1 interaction were shown next to the legend.

## Discussion

Here, we report the relationships between respiratory activity and mitochondrial MDH1-CIT1 multienzyme complex interaction in living yeast cells. MDH1-CIT1 multienzyme complex dynamically associated and dissociated in response to respiratory status. This study is the first in vivo evidence in any organism showing the dynamic association and dissociation of the TCA cycle metabolon in real time. The MDH-CS metabolon, specifically, is considered essential for the forward TCA cycle flux due to its ability to overcome the unfavorable thermodynamics of MDH reaction under physiological oxaloacetate concentrations (Beeckmans and Kanarek, 1981; Huang et al., 2018). The results of this study demonstrate that the MDH1-CIT1 multienzyme complex associates when the TCA cycle is active and dissociates when it is suppressed, supporting the idea that metabolon dynamics are related to pathway regulation. Given that the MDH-CS metabolon is proposed to enhance pathway reactions (Bulutoglu et al., 2016; Fukuda et al., 2008), its dynamic assembly and disassembly would influence pathway activity. Together, these observations align with the proposed theory that metabolon dynamics act as a regulatory mechanism of metabolic flux (Noor et al., 2014; Obata, 2020).

The MDH1-CIT1 complex dissociated under conditions suppressing the TCA cycle. TCA cycle inhibition by arsenite and AOA induced dissociation of the MDH1-CIT1 complex (Figure 3B&C). In a physiological condition, the MDH1-CIT1 complex dissociated when Crabtree-inducing fermentable sugars were supplied to the media (Figures 1, 2, and Figure 2 – figure supplement 1-3). The Crabtree effect redistributes respiratory flux to stimulate fermentation and repress aerobic respiratory pathways, including the TCA cycle (Gancedo, 1998; Ronne, 1995). The metabolic shift by the Crabtree effect is a concerted action of multiple mechanisms, such as the suppression of enzyme gene expressions and the inhibition of mitochondrial transporters (Ahmadzadeh et al., 1996). Interestingly, glucose-induced rapid repression of mitochondrial respiration occurs at a rate that is too great to be explained only by the inhibition of enzyme synthesis, and additional forms of regulation are anticipated to be involved (Ahmadzadeh et al., 1996). The dissociation of the TCA cycle metabolon coincided with this repression (Figure 2A), highlighting metabolon dynamics as a potential additional mechanism to swiftly downregulate mitochondrial respiration during the Crabtree effect.

On the other hand, the MDH1-CIT1 complex interaction was enhanced under conditions that activate the TCA cycle. An increase in TCA cycle activity by the addition of acetate (Orlandi et al., 2014; Vilela-Moura et al., 2011) enhanced the MDH1-CIT1 complex interaction (Figure 3A), suggesting the role of the MDH-CS metabolon in facilitating TCA cycle activity.

However, ETC inhibitions also increased MDH1-CIT1 interaction (Figure 4A-C), despite their inhibitory effects on oxidative respiration (Figure 4 - figure supplement 1) and the TCA cycle. This apparent contradiction can be explained by the transient nature of the response. The rise in MDH1–CIT1 interaction following ETC inhibition was transient, suggesting that it does not represent steady-state changes in respiratory activity. Instead, it is more likely linked to acute changes in the mitochondrial matrix microenvironment. Notably, the timing of the increased interaction coincided with changes in matrix pH, supporting a role for pH-dependent regulation (see below). These findings align with our model, which posits that the MDH1–CIT1 interaction is controlled by microenvironmental changes accompanying shifts in respiration. In the case of ETC inhibition, transient perturbations of the matrix microenvironment appear to be disconnected from steady-state respiratory output, thereby promoting MDH1–CIT1 interaction despite overall inhibition of respiration.

We also investigated the molecular mechanisms regulating MDH1-CIT1 complex dynamics. We previously reported that protein conformation changes induced by the solution pH and allosteric regulators affect porcine MDH-CS complex interaction in vitro (Omini et al., 2021). The recombinant yeast MDH1 and CIT1 showed similar responses to the porcine enzymes; MDH1-CIT1 affinity was higher in low pH and in the presence of upstream TCA cycle intermediates (malate and fumarate) but disrupted in the presence of reaction product (citrate; Figure 5). These factors can affect the TCA cycle metabolon formation in vivo. The relationship between these factors and the MDH1-CIT1 interaction observed in this study is summarized in Figure 6A.

**Figure 6.**
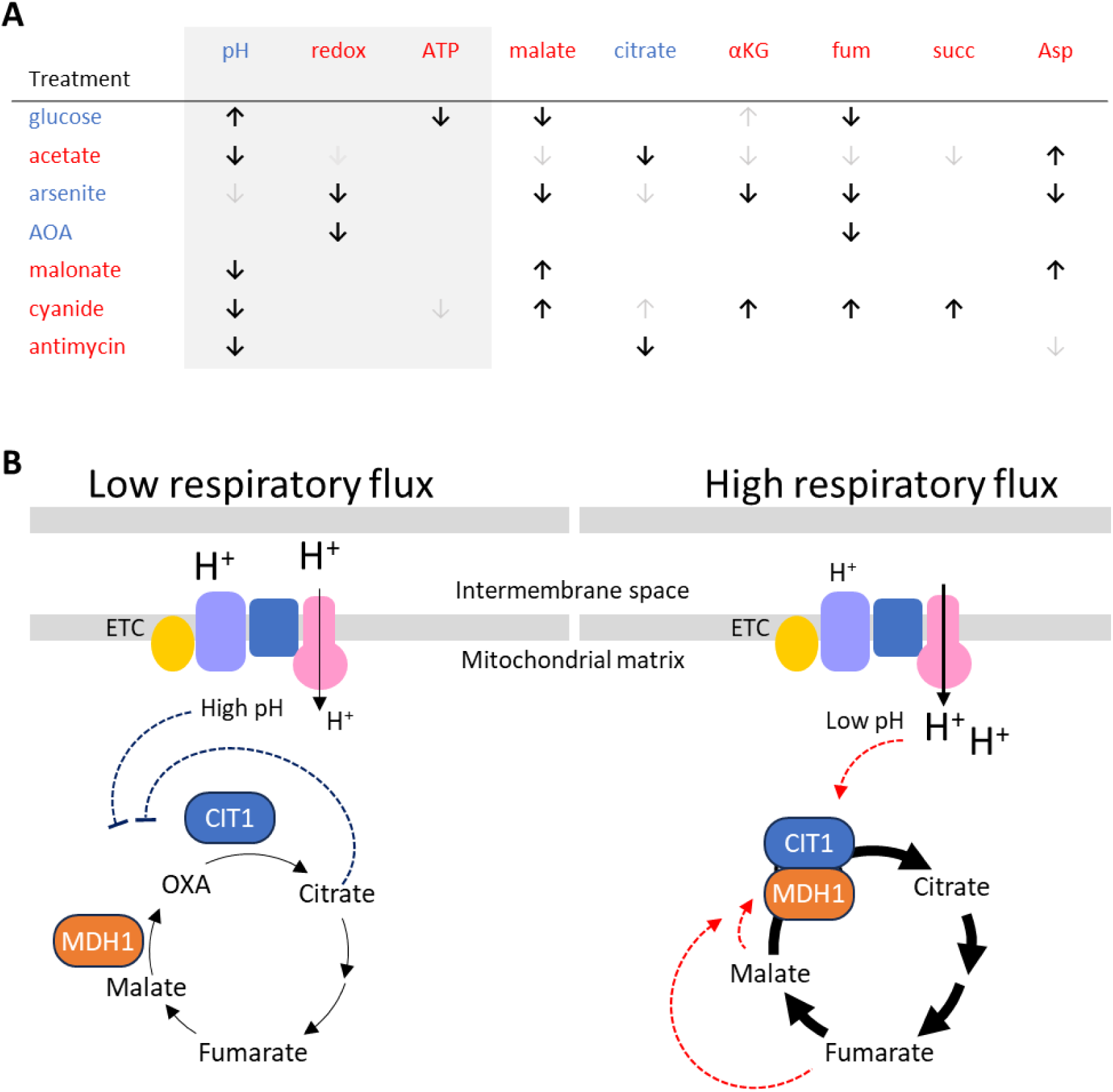
Relationship between the respiratory metabolism and the MDH1-CIT1 metabolon association. **(A)** Summary of the effects of metabolic treatments on MDH1-CIT1 complex association, mitochondrial matrix parameters (pH, redox state, and ATP levels), and cellular metabolite levels. Each row corresponds to a specific metabolic perturbation. Blue and red labels indicate treatments that decrease or increase the complex association, respectively. Upward and downward arrows indicate increases and decreases in each parameter, respectively. Bold arrows denote Growth rate and enzyme activity of NanoBiT reporter changes that are consistent with the observed in vivo alterations in MDH1–CIT1 interaction. The color of the parameter and metabolite indicates its effect on MDH1-CIT1 interaction in vitro (blue, inhibitory; red, promotive). αKG, α-ketoglutarate; fum, fumarate; succ, succinate. **(B)** A diagram depicting the proposed regulatory mechanism of the MDH1-CIT1 metabolon association. In conditions with low respiratory flux, the MDH1-CIT1 multienzyme complex dissociates, and the TCA cycle flux reduces. Reduced ETC flux results in higher mitochondrial matrix pH, which reduces MDH1-CIT1 affinity. When the respiratory flux and the TCA cycle flux are high, MDH1-CIT1 metabolon associates and likely channels the intermediate oxaloacetate (OXA). High ETC flux lowers mitochondrial matrix pH and enhances the MDH1-CIT1 interaction. The TCA cycle intermediates affect MDH1-CIT1 metabolon formation; fumarate and malate enhance (red arrows with dotted lines), and citrate inhibits (blue arrows with dotted lines) the interaction. The arrow thickness represents the metabolic fluxes.

We observed an enhanced MDH1-CIT1 interaction under conditions that lower mitochondrial matrix pH in vivo. The mitochondrial matrix pH decreased after adding acetate, corresponding to the enhanced MDH1-CIT1 interaction (Figure 3 A, D). The time courses of the temporal increase in MDH1-CIT1 interaction and pH decrease were inversely related following the complex III and IV inhibitions (Figure 4B, C, E, F; Figure 6A). The decrease in the pH from 7.5 to 6.0 ranges is remarkable since the *K*_d_ of MDH1-CIT1 association decreases by an order of magnitude within this pH range (Figure 5A). The mitochondrial matrix acidification under respiratory conditions likely favors the MDH1-CIT1 interaction. On the other hand, the mitochondria matrix pH was maintained around 7.2 in the presence of glucose (Figure 2D), which likely weakens the MDH1-CIT1 complex interaction. These results indicate that mitochondrial matrix pH, which is related to proton transport activity by the ETC, can stabilize or destabilize the MDH1-CIT1 complex, thereby regulating the MDH1-CIT1 interaction in response to respiratory activity (Figure 6B). Enhancing the MDH1-CIT1 interaction with the uncoupler CCCP (Figure 4 – figure supplement) also supports the prominent effect of pH.

Nevertheless, mitochondrial pH may influence the MDH1-CIT1 association, but it is not always the predominant factor regulating the interaction. The observed changes in pH and interaction do not consistently occur simultaneously following the Crabtree induction and acetate supplementation (Figs 2 & 3). TCA cycle inhibitions reduce the MDH1-CIT1 interaction without significantly affecting mitochondrial matrix pH (Figure 3B, C, E, F). These findings suggest that mitochondrial matrix pH can be a contributing factor, either favoring or disfavoring the interaction, rather than the primary regulator, which appears to be the direct induction and dissociation of the MDH1-CIT1 complex.

Little evidence in this study supports a primary role for mitochondrial matrix redox state or ATP levels in controlling MDH1–CIT1 interaction. Our in vitro analyses demonstrate that NAD(H) and ATP can modulate MDH1–CIT1 interaction (Figure 5 - figure supplement 1B; Omini et al., 2021), suggesting that these factors may contribute to complex dynamics. Although the lower matrix ATP level following glucose supplementation (Figure 2D) and matrix reduction upon arsenite and AOA-induced TCA cycle inhibition (Figure 3H&I) associated with decreased interaction (Figure 2A and 3B&C), these relationships were not consistently observed across conditions (Figure 6A), indicating that neither redox state nor ATP levels predict complex association in vivo. However, we cannot rule out their contribution under specific metabolic contexts not captured in the current study.

The TCA cycle intermediates and cofactors can also regulate the MDH1-CIT1 interaction, considering their effects on the interaction in vitro (Omini et al., 2021). The yeast enzymes showed responses to malate, α-ketoglutarate, succinate, and citrate (Figure 5B, Figure 5 – figure supplement 1) similar to the enzymes of other organisms (Omini et al., 2021; Tompa et al., 1987; Wu et al., 2015), while fumarate is newly identified as an effector of the MDH-CS interaction.

Especially, fumarate, malate, and citrate showed significant influences on yeast enzymes (Figure 5B). Cellular levels of these effector metabolites significantly altered under the conditions tested in this study (Figure 2E, 3M, 4M, 6A). Increased and decreased levels of malate and fumarate following the TCA cycle and ETC inhibitions (Figure 3M&4M), respectively, are likely related to the MDH1-CIT1 interaction since malate enhances the interaction (Figure 5D; Omini et al., 2021). These results indicate the involvement of metabolite effectors, such as malate and fumarate, in the regulation of the MDH1-CIT1 interaction (Figure 6). However, their precise effects must be evaluated through site-specific, time-dependent perturbation and quantification of metabolite levels in the mitochondrial matrix, as a whole-cell metabolite profile may not reflect the metabolite concentrations that directly influence the complex association.

Aside from MDH1-CIT1 interaction dynamics, our results highlight the complex regulation of TCA cycle metabolism. The activation and inhibition of respiratory activity resulted in diverse metabolic phenotypes (Figure 6A, Supplementary Dataset 1), where intermediate levels did not simply reflect overall pathway activity. This complexity stems from the distinct mechanisms of each inhibitor, such as arsenite affecting α-ketoglutarate dehydrogenase and AOA disrupting the malate-aspartate shuttle (Cavero et al., 2003; Eto et al., 1999; Lee et al., 2011; Liu and Butow, 2006), off-target effects of the inhibitors, and from the adaptive reorganization of intersecting metabolic networks to bypass local blockades (Herrgård et al., 2008; Lehtinen et al., 2013; Rogers et al., 2021). These diverse metabolic phenotypes allow us to assess the relationships between metabolites and metabolon assembly independently of respiratory activity.

This study demonstrates that the TCA cycle MDH1-CIT1 multienzyme complex dynamically interacts in response to the cellular respiratory status. Cues of cellular respiratory state may be transmitted to the multienzyme complex, at least in part, by mitochondrial matrix microenvironment and metabolite levels. In particular, mitochondrial matrix pH and malate and fumarate levels likely have significant effects on the stability of the MDH1-CIT1 complex, since these factors strongly influence complex affinity, and their in vivo changes coincide with complex dynamics (Figure 6). However, none of these factors consistently correlates with the MDH1-CIT1 interaction (Figure 6A), suggesting that none of them is a predominant regulator, but multiple factors work together in a cooperative manner to regulate the formation of the multienzyme complex. This coordinated regulation probably serves to fine-tune the flux of the TCA cycle, allowing it to adapt efficiently to varying metabolic demands and maintain cellular homeostasis. Although we focused on allosteric regulators in this study, further factors are potentially involved in the MDH1-CIT1 complex regulation. For example, 44 and 33 post-translational modifications have been identified in CIT1 and MDH1, respectively (Bhagwat et al., 2021; Henriksen et al., 2012; Holt et al., 2009; Lanz et al., 2021; Reinders et al., 2007; Swaney et al., 2013; Weinert et al., 2013), some of which likely affect the MDH1-CIT1 complex affinity. Various scaffolding molecules, such as long noncoding RNAs, lipid layers, and scaffolding proteins, have been shown to stabilize the multienzyme complexes in other systems (Zhu et al., 2022). Future studies should investigate the effects of these factors to understand the regulatory mechanisms for MDH1-CIT1 interaction.

Changes in environmental conditions and nutrient availability occur swiftly, and a corresponding change in metabolic flux must occur at a similar rate for the successful adaptation of living cells. Dynamic metabolons can be a system to regulate and fine-tune metabolic network flux quickly, allowing cells to maintain metabolic homeostasis in rapidly fluctuating environments (Obata, 2019). Considering the thermodynamic unfavorability of the forward MDH reaction and that substrate channeling overcomes this thermodynamic barrier in vitro, the MDH1-CIT1 complex formation likely enhances the forward TCA cycle flux (Bulutoglu et al., 2016; Fukuda et al., 2008; Sweetlove et al., 2010). The dynamic assembly of the metabolon probably provides fine-tuning of TCA cycle activity in response to respiratory demand. While the present study delineates the environmental and metabolic factors governing this dynamic assembly, future studies utilizing targeted genetic tools to independently modulate specific variables will be critical for dissecting the contributions of individual regulatory factors in vivo. Further studies assessing the impacts of metabolon formation on metabolic pathway flux in living cells are essential to understand the functions of metabolons in metabolic network regulation. Elucidating the regulatory system of the TCA cycle metabolon can lead to a novel strategy to manipulate the TCA cycle flux in metabolic engineering to achieve efficient industrial production of various molecules requiring TCA cycle intermediates as substrates or to control the Warburg effect to suppress cancer cells.

## Materials and Methods

### Strains, media, and culture conditions

*Saccharomyces cerevisiae* BY4741 (MATa his3*Δ*1 leu2*Δ*0 met15*Δ*0 ura3*Δ*0) was used as the background strain. Cells were grown in synthetic complete medium (SD) containing 0.67% yeast nitrogen base lacking amino acids (Research Products International, Mt. Prospect, IL, USA) with 2% w/v D-raffinose and 1% amino acid complete drop-out mix. The complete amino acid drop-out mix was replaced with the amino acid drop-out mix lacking leucine for GoAteam expressing cells or the amino acid drop-out mix lacking uracil for RoGFP and pHluorin expressing cells. Cell growth cultures were incubated with shaking at 220 rpm at 28°C in an incubator shaker. Cell cultures were grown to exponential phase with an OD_600_ = 0.5 to 1.0 in all analyses.

## Oxygen consumption rate measurement

Oxygen consumption was measured using a Clark-type electrode (Oxygraph, Hansatech Instruments, Norfolk, UK) as described previously (Agrimi et al., 2011). Cells were grown to exponential phase OD_600_=0.5, harvested, centrifuged at 3,000 x *g* for 5 min at 4°C, and resuspended in growth medium to obtain a density of OD_600_=5.0. A total of 1 ml reaction volume consisting of equal parts of 445 µl cell culture, 445 µl growth media, and 10 µl of each inhibitor was added to the oxygraph chamber. Sodium malonate (20 mM), antimycin A (10 µM), sodium cyanide (0.5 mM), and oligomycin (1 mM) were used as specific inhibitors of electron transport chain complex II, III, IV, and V, respectively. The change in oxygen concentration was followed subsequently. Oxygen consumption rates were determined before and after the addition of each inhibitor from the slope of a plot of O_2_ concentration versus time. All measurements were conducted in triplicates.

### Generation of the *Saccharomyces cerevisiae* reporter strains with tagged MDH1 and CIT1

The NanoBiT split NanoLUC luciferase complementation system (Dixon et al., 2016) was adopted to monitor MDH1-CIT1 interaction in yeast cells. The yeast lines expressing MDH1 and CIT1 proteins fused with the small (SmBiT) and large (LgBiT) NanoBiT subunits, respectively, were generated by inserting the yeast codon-optimized tag-coding sequences into the BY4741 genome following the scarless C-terminal tagging procedure (Landgraf et al., 2016). The full-length NanoLUC, LgBiT, and SmBiT coding sequences were integrated to direct downstream of the Mdh1 (chrIV:3300230) and Cit1 (chrX:303993) genes on the BY4741 genome for C-terminal fusion. The LgBiT and SmBiT sequences include flexible linkers (ACKIPNDLKQKVMNH; (Hu et al., 2002)) with HA and cMyc epitope sequences, respectively, for immunological detections. Details of the vector construction procedure, primers, and vectors are described in the Supplementary Method.

Briefly, we generated the plasmids with integration cassettes composed of the linker, 5’ half of tag-coding sequence, URA3 selection marker, and 3’ half of tag-coding sequence in this order using the Golden Braid technology (Sarrion-Perdigones et al., 2013). The halves of the tag-coding sequences have an overlapping sequence. The integration cassettes were amplified by PCR using the primers with around 40 bases 5’ extension with sequences homologous to the franking region of the tag insertion sites. BY4741 cells were transformed with the purified PCR products to insert the cassettes into the target sites directly downstream of the Mdh1 and Cit1 genes by homologous recombination due to the 5’ extension of the primers. The transformants were selected on uracil-deficient plates. To pop-out the URA3 marker and reconstruct NanoLUC and NanoBiT subunits, the transformants were cultured overnight in uracil-containing YPD media. The URA3 gene was excluded from the genome by recombination based on the overlapping sequences in the cassette. The cell suspension was spread on SD plates containing 1 mg ml^-1^ 5-fluoroorotic acid (5-FOA) to select the cell lines without the URA3 gene. SmBiT was initially fused with the Mdh1 gene, and the resulting strain was further transformed to fuse LgBiT with Cit1 (MDH1/CIT1-BiT strain). Another strain expressing MDH1 fused with full-length NanoLUC luciferase was also generated following the same procedure to monitor MDH1 protein levels (MDH1-nLUC strain).

## In vivo MDH1-CIT1 interaction measurement

The MDH1/CIT1-BiT strain expressing MDH1 and CIT1 enzymes tagged with split halves of luciferase enzyme was grown to OD_600_ = 0.35-0.45. Cells were collected and resuspended to obtain a cell density of OD_600_=2.0 in fresh SD-Raff media. Each sample consisted of 80 µl of media, 10 µl of cells, and 10 µl of 50x furimazine luciferase substrate (Promega, Madison, WI, USA). The luminescence signal was measured every five min with a microplate reader (CLARIOSTAR Plus, BMG LABTECH, Ortenberg, Germany) at 28°C with 200 rpm shaking at the beginning of each cycle. Baseline luminescence was measured for 20 min before the treatment was applied. The time-dependent luminescence was measured for an additional 80 min after treatment. Relative luminescence Unit (RLU) was calculated by normalizing the luciferase signals by the average signals during three pre-treatment time points. The protein levels of MDH1 and CIT1 were assessed using yeast lines expressing these proteins tagged with full-length NanoLUC luciferase. Using the luminescence data of these relative protein levels, we calculated normalized interaction index by dividing the NanoBiT interaction signal by the product of the relative abundances of both proteins:

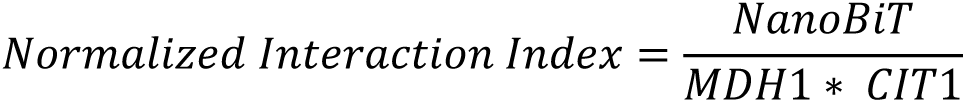

In this formula, NanoBiT, MDH1, and CIT1 are the relative luminescence levels at each time point.

## Western blotting

Cells were grown to OD_600_=0.5 and cell pellet was collected. Cells were first pretreated with 2 M LiAc and 0.4 M NaOH to permeabilize the cells and then treated with SDS-PAGE sample buffer to extract proteins according to the method described by Zhang et al. (Zhang et al., 2011). The cell lysate was centrifuged at 27,000 x *g* for 10 min at 4°C and the proteins were detected by SDS-PAGE and western blotting following the method described previously (Rajasekaran et al., 2022). cMyc Tag Monoclonal Antibody (MA1213161MG, Thermo Scientific) and HA Tag Monoclonal Antibody (LSG26183, Thermo Fisher Scientific) were used as primary antibodies to detect MDH1 and CIT1, respectively. DLD1 antibody generated by S. Claypool laboratory at the Johns Hopkins University (Baile et al., 2013), and PGK1 Monoclonal Antibody (Invitrogen 459250, Thermo Fisher Scientific) was used to detect phosphoglycerate kinase as a housekeeping protein and internal standard.

## In vitro MDH1-CIT1 interaction measurement

The recombinant MDH1 and CIT1 were produced accordingsuw to the method described in the Supplementary Methods. The interaction between recombinant MDH1 and CIT1 enzymes were analyzed by microscale thermophoresis (MST) according to the method described by Omini et al. (Omini et al., 2021) with slight modifications. Base MST buffer contained 50 mM Tris-HCL (pH 8), 150 mM NaCl, 10 mM MgCl_2_, 5 mM DTT, and 0.05% Tween-20. Recombinant MDH1 (10 µM) was labeled with Protein Labeling Kit RED-NHS 2nd Generation (NanoTemper, München, Germany) and used as the target. CIT1 was used as the ligand. Two-times serial dilution of 80 µM CIT1 was conducted for 16 concentrations. A total of 10 µL of CIT1, 10 µL MST buffer, and 10 µL labeled MDH1 were mixed and loaded to Monolith NT.115 Capillaries (NanoTemper).

Capillaries were incubated at room temperature for 1 min, and the interaction was analyzed by MST using Monolith NT.115 (NanoTemper). To test the effect of pH on MDH1-CIT1 interaction, 50 mM tris buffers with pH 6.0, 7.5, and 8.0 were prepared and used as the MST buffer for sample preparation. To test the effect of reducing environment on interaction, various concentrations (5 mM, 2.5 mM, 1.25 mM) of DTT were prepared with the MST buffer and used for sample preparation. To test the effect of ATP level on interaction, an MST buffer containing different concentrations of ATP (5 mM, 2.5 mM, 1.25 mM) was made and used for sample preparation. To test the effect of metabolite availability on the interaction, MST buffer containing respective metabolites at 10 mM concentration was made and used for sample preparation and experiments.

## Enzyme activity assays

Cells were grown to an exponential phase with OD_600_ of 0.5, and 2 ml of the cells were harvested for enzyme activity assay. Yeast cell lysates were prepared by disruption with glass beads as described previously (Mukherjee et al., 2020), with the lysis buffer omitted butylhydroxytoluene. The protein concentration of the lysate was determined using Pierce™ BCA Protein Assay Kit (Thermo Fischer Scientific). CS activity was determined using a method described previously (Srere, 1969) that measures free thiols by coupling the citrate synthase reaction to thiol reaction with 5, 5’-dithiobis-(2-nitrobenzoate) (DTNB). The CS enzyme activity assay mixture contained 154 µL of distilled water, 20 µL of 1 mM DTNB, and 6 µL of 10 mM acetyl CoA. The citrate synthase reaction was initiated by the addition of 10 µL of 10 mM oxaloacetate. The absorption at 412 nm was followed to measure citrate synthase activity. MDH activity assay mixture contained 50 mM TES (pH 7.2), 5 mM MgCl_2_, 0.2 mM NADH, and 0.05% Triton X100. To obtain a total reaction volume of 300 µL, 285 µL of the assay mixture and 5 µL cell lysate were added to the wells, and the reaction was initiated with 10 µL of 30 mM oxaloacetate. Reduction of NADH was followed at 340 nm to determine MDH enzyme activity. Enzyme activity was measured using a microplate reader absorbance function (CLARIOSTAR Plus, BMG LABTECH).

## Metabolite profiling

Yeast cells were grown to the exponential phase (OD_600_∼0.5), and treatments were applied and incubated for the time described in the figure legends. Cellular metabolites were extracted following the protocol described by Obata et al. (Obata et al., 2013). The cells in 1 mL culture were harvested by vacuum filtration using a membrane filter (0.45 µm HV Durapore 25 mm diameter; MilliporeSigma, Burlington, MA, USA). The filter was put into a 2 mL microcentrifuge tube, flash-frozen in liquid nitrogen, and stored at −80°C. The metabolites were extracted from the filtered cells with methanol: water: chloroform, and 50 µL aliquot was dried down by vacuum centrifugation. Dried metabolites were derivatized with methoxyamine hydrochloride in pyridine and further trimethylsylilated by N-Methyl-N-(trimethylsilyl) trifluoroacetamide (MilliporeSigma). Derivatized samples were analyzed by 7200 GC-QTOF system (Agilent, Santa Clara, CA, USA) exactly as described in Wase et al. (Wase et al., 2022). Each metabolite’s peak height was normalized by the peak height of the internal standard (ribitol) to represent relative levels of metabolite.

## Expression of mitochondrial biosensors

The pH, redox state, and ATP levels in the mitochondrial matrix were measured using the mito-Go Ateam2, mito-ROGFP1, and pHluorin (pAG416-COX4-pHluorin, URA selection marker) fluorescence biosensors, respectively, specifically localizing in the mitochondrial matrix. The mito-Go Ateam2 (p415-GPDpro-mito GO ATeam) and mito-roGFP1 (p416-GPDpro-mito roGFP) encoding plasmids (Vevea et al., 2013) were generous gifts from Dr. Liza Pon at Department of Pathology and Cell Biology, Columbia University. The pHluorin encoded plasmid (pAG416-COX4-pHluorin;(Ayer et al., 2013)) was a generous gift from Dr. Anita Ayer at the University of New South Wales, Sydney, Australia. The plasmids encoding these biosensors were transformed into split-Luc tagged MDH1/CIT1 strain using the lithium acetate method (Chen et al., 1992). The p416-GPDpro-mito roGFP and pAG416-COX4-pHluorin harboring cells were selected on URA- media, and the cells with p415-GPDpro-mito GO ATeam were selected on Leu- media.

The localization of biosensors within the mitochondria was confirmed by confocal microscopy. Cells were grown in SD selection media to OD_600_=0.50. Cells were stained with MitoTracker Deep Red FM (ThermoFisher Scientific) in a 100 nM dye solution for 30 min with shaking at 28 °C to visualize the mitochondria. The stained cells were resuspended in fresh 10 mM HEPES buffer (pH 7.4) with 2% raffinose to obtain a final concentration of OD_600_=10. Confocal imaging was performed on an A1R-Ti2 Confocal Laser Scanning Microscope (Nikon, Tokyo, Japan) with a Plan Apo 60x 1.40 Oil lens 0.17 WD 0.13 (Nikon) and a 2x digital zoom for a total magnification of 1200x. Brightfield and fluorescent images were captured. Imaging was done sequentially with excitation at 405 nm and emission at 425-475 nm for pHluorin and mito-roGFP1, excitation at 488 nm and emission at 500-550 nm for mito-GoAteam2, and excitation at 640 nm and emission at 663-738 nm for MitoTracker. Images were collected with NIS Elements software (Nikon) and processed with ImageJ analysis software (Vevea et al., 2013).

## Measurement of redox state, pH, and ATP level in the mitochondrial matrix

Cells expressing pHluorin, mito-roGFP1, and mito-GoAteam2 were grown to OD_600_=0.50 in their respective selection media. Cells were prepared for fluorescence measurement according to the method described by Morgan *et al* (Morgan et al., 2011). Time-based fluorescence intensity was measured using a microplate reader (CLARIOSTAR Plus, BMG LABTECH) at 28°C with shaking at 200 rpm with 20 min of baseline measurement before treatments. In all experiments, the strains harboring empty vectors were grown simultaneously as a reference for background fluorescence at the different excitation wavelengths. Background fluorescence was subtracted from fluorescence intensity from cells expressing biosensors.

For the pH measurement, the ratio of emission intensity at 510 nm resulting from excitation of pHluorin at 390 nm and 470 nm was calculated (R390/470) using pHluorin expressing strain. The mitochondrial matrix pH was calculated from the R390/470 ratio using a calibration curve generated according to the method described by Orij *et al* (Orij et al., 2009). Briefly, pHluorin-expressing cells were permeabilized by resuspension in PBS containing 100 µg ml-1 digitonin for 10 min and washed with PBS. Cells were resuspended in citric acid/Na2HPO4 buffer of pH values ranging from 5.0 to 9.0. The ratio of pHluorin emission at 510 nm upon excitation at 390 and 470 nm (R390/470) was plotted against buffer pH to obtain a calibration curve.

The mitochondrial matrix redox state was measured using mito-roGFP1 expressing strain. The ratio of emission intensity at 510 nm from excitation at 365 nm and 470 nm (R470/365) was measured. Mito-roGFP1 in situ calibration was performed following the method described by (Vevea et al., 2013). Digitonin-treated mito-roGFP1 expressing cells were incubated in 0, 5, and 10 mM H_2_O_2_ and DTT for 20 min at 28°C with shaking at 200 rpm. The R470/365 was plotted against the redox potential of the solutions calculated using the formula described previously (Morgan et al., 2011).

Relative ATP level was determined by measuring the ratio of the emission intensity of mito-GoAteam2 at 510 nm and 560 nm with excitation at 470 nm (R560/510). The ATP levels were analyzed as the relative value without calibration to absolute concentrations.

## Statistical Analysis

The differences between the control and test samples were evaluated by a two-tailed unpaired Student’s *t*-test. *p* < 0.05 was considered a statistically significant difference. For the time course data, the test was applied at each time point. All data was obtained from triplicated independent experiments.

## Acknowledgments

This study is supported by the NSF CAREER Award to TO under Grant No. 1845451. Dr. Liza Pon at Columbia University kindly gifted us the mito-Go Ateam2 and mito-ROGFP1 biosensors, and Dr. Anita Ayer at the University of New South Wales, Sydney, Australia, gifted us the pHluorin mitochondrial biosensor. Dr. Oleh Khalimonchuk at the University of Nebraska-Lincoln provided us with the anti-DLD1 antibody. The Microscopy work assisted by Terri Fangman was carried out at the Microscopy Research Core Facility of the Center for Biotechnology at the University of Nebraska-Lincoln, which is partially funded by the Nebraska Center for Integrated Biomolecular Communication COBRE grant (P20 GM113126 and NIGMS) and the Nebraska Research Initiative. This paper is based on a dissertation submitted by Joy Omini to fulfill the requirements for the degree of Doctor of Philosophy, University of Nebraska-Lincoln (Omini, 2024).

---

**Figure 1 – figure supplement 1.**
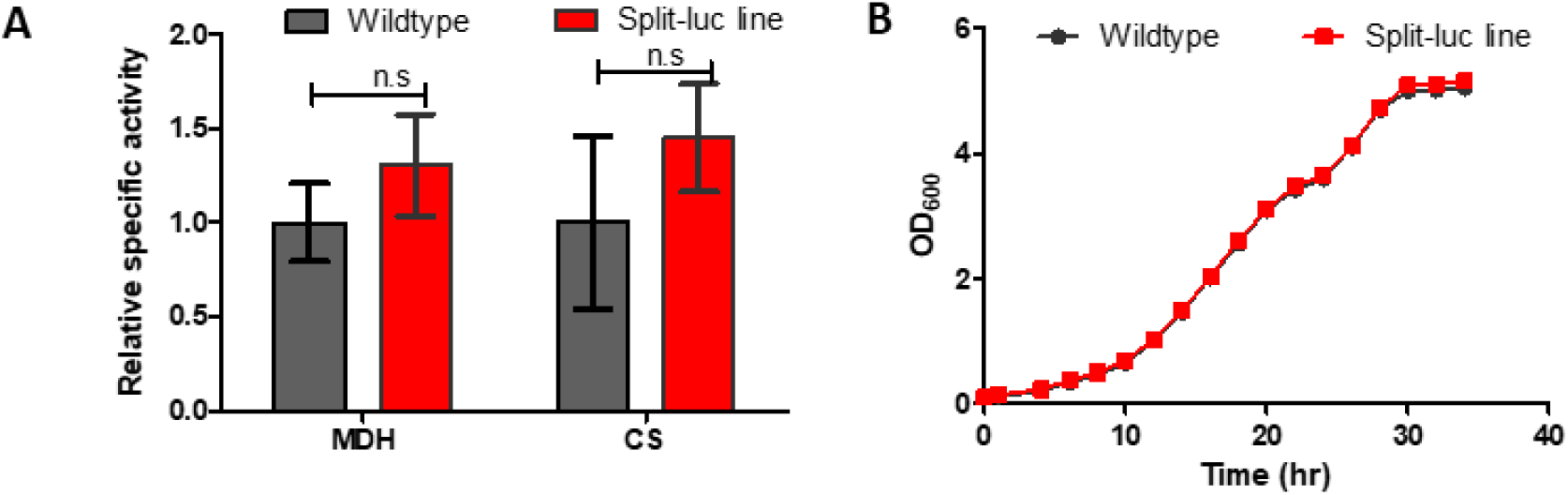
Growth rate and enzyme activity of NanoBiT reporter strain. (A) Extractable cellular MDH and CS enzyme activities in the wildtype (black) and NanoBiT reporter strain (Split-luc line, red). (B) Growth of cells in SD-raff media monitored as culture OD600. Data is presented as mean ± s.d. Statistical differences against the wildtype samples were assessed by Student’s *t*-test at each time point. n.s., not significant (*p*>0.05).

**Figure 1 – figure supplement 2.**
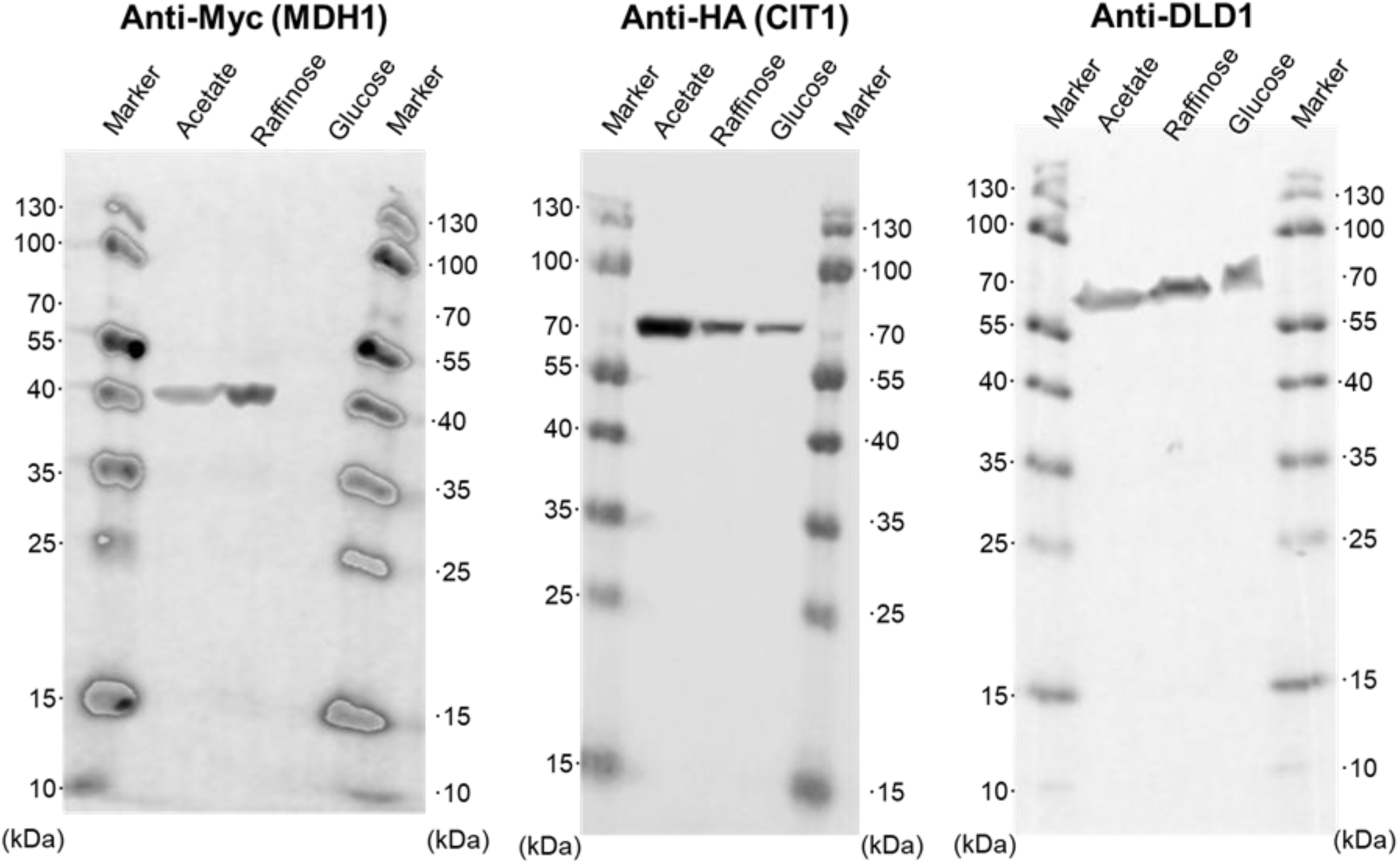
Full Western blotting image showing MDH1 and CIT1 protein abundance in. Figure 1C. The numbers on the side of the images indicate the size of the molecular weight markers (kDa).

**Figure 2 – figure supplement 1.**
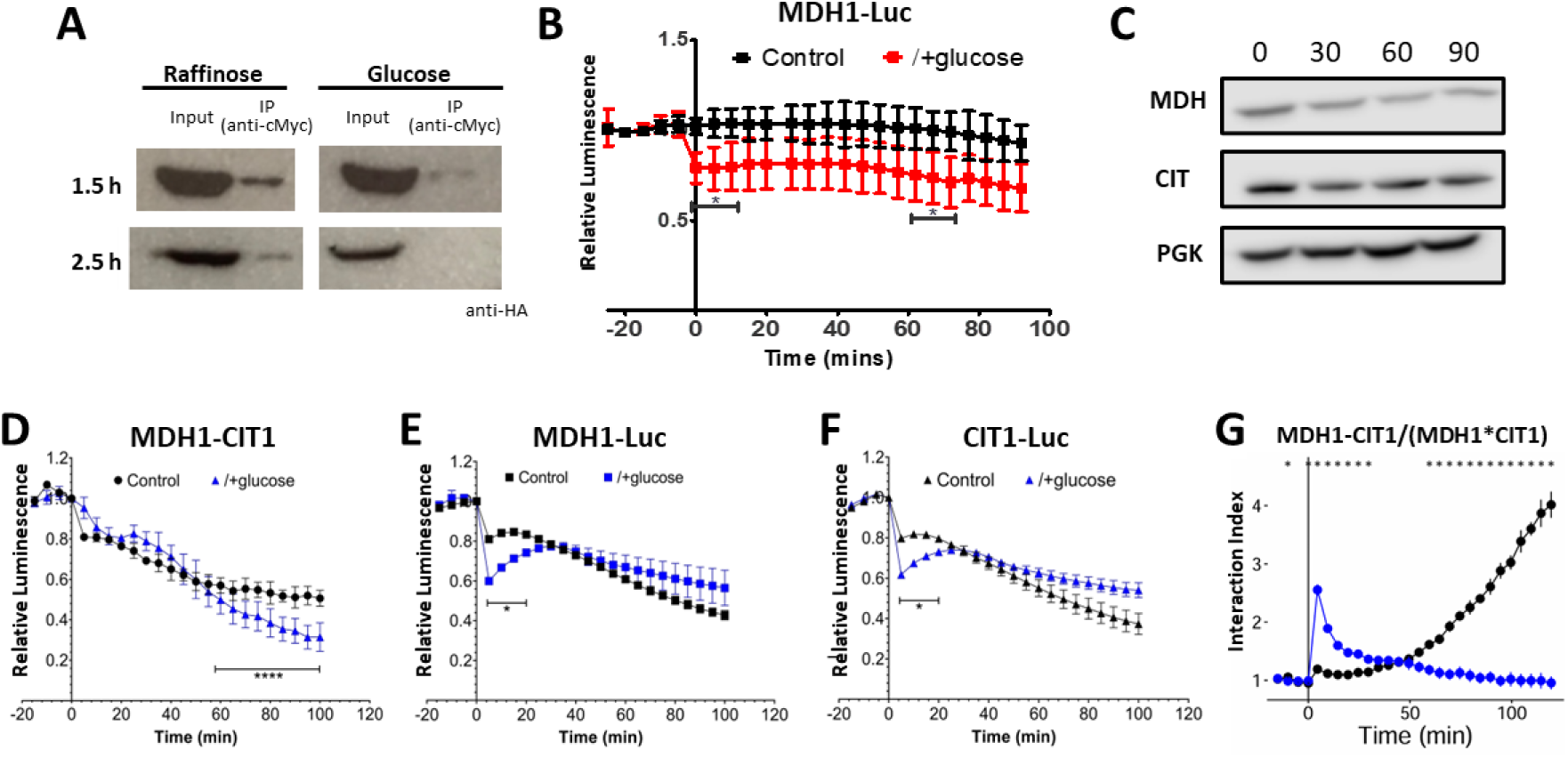
MDH1-CIT1 interaction and protein abundance following the application of glucose. **(A)** Co-immunoprecipitation of CIT1 with MDH1. In the Split-luc line, MDH1-LgBiT and CIT1-SmBiT harbor cMyc and HA tag in the linker sequence. Proteins were extracted from the Split-luc line grown in SD-faff media at 1.5 and 2.5 hours following 2% glucose application. MDH1 was precipitated with anti-cMyc agarose and co-precipitated CIT1 was detected by anti-HA antibody following SDS-PAGE separation. **(B)** MDH1 protein levels monitored by the luminescence of MDH1 fused with full-length NanoLUC luciferase. **(C)** Western blot analysis of MDH1 and CIT1 protein levels after 0, 30, 60, and 90 min of Crabtree induction. Phosphoglycerate kinase (PGK) was detected as a loading control. **(D)** NanoBIT signal indicating MDH1-CIT1 interaction in a repeated glucose supplementation experiment. SD-Raff-grown cells were treated with 2% glucose at 0 min (blue). **(E)** MDH1 protein levels monitored by the luminescence of MDH1 fused with full-length nanoLUC luciferase in the repeated experiment. **(F)** MDH1 protein levels monitored by the luminescence of CIT1 fused with full-length nanoLUC luciferase in the repeated experiment. Relative luminescence was calculated by normalizing the luciferase signals by the average signals during three pre-treatment time points. Statistical differences against the control samples were assessed by Student’s *t*-test at each time point (N=4). Asterisks indicate significant difference (*, *p*<0.05; ****, *p*<0.0001; ns, not significant). **(G)** Interaction index calculated by normalizing NanoBiT signal by those of MDH1-nanoLUC and CIT1-nanoLUC. Asterisks indicate time points with statistically significant differences (p<0.05) between control and glucose conditions (N=4).

**Figure 2 – figure supplement 2.**
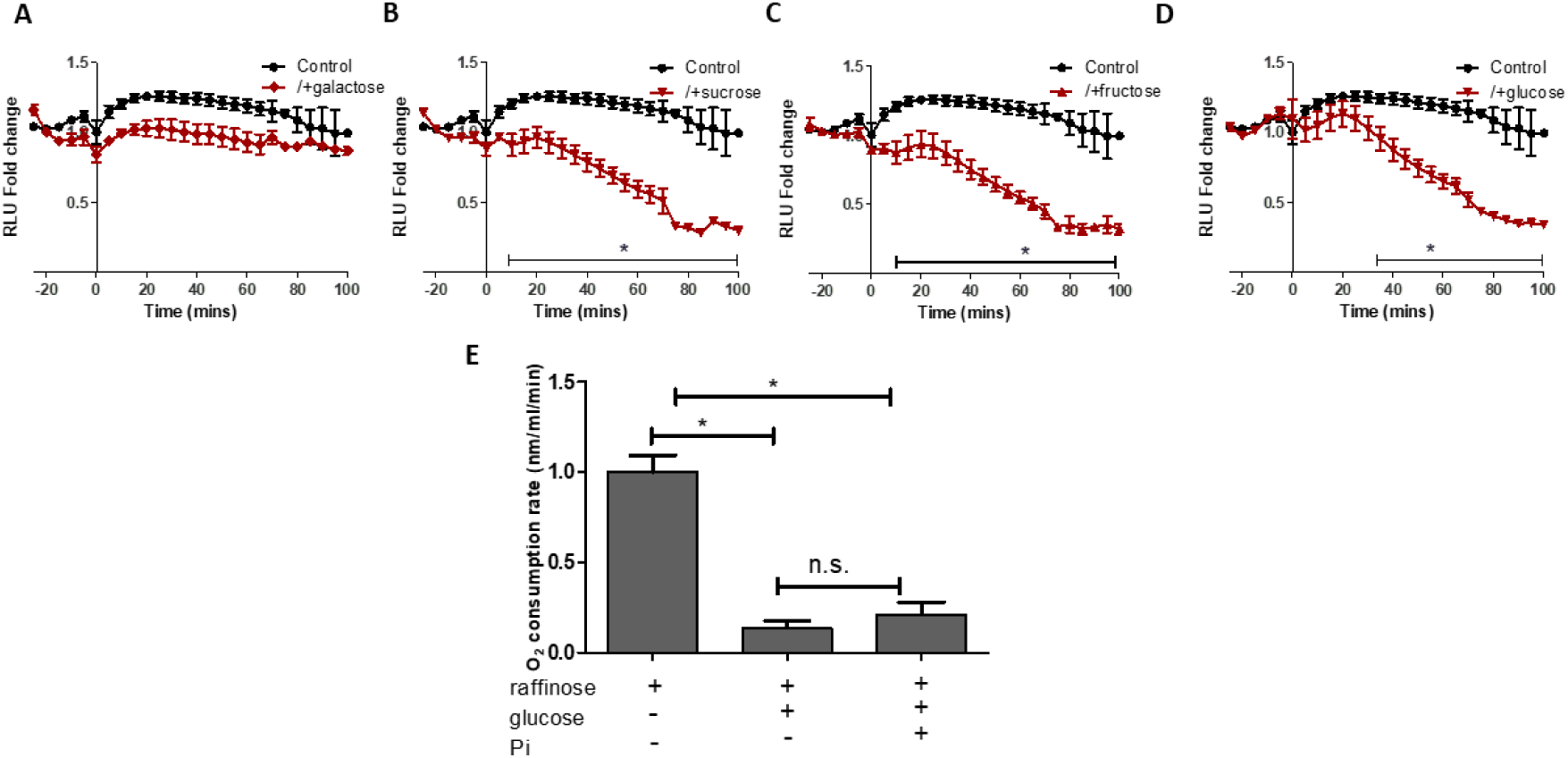
Effects of sugars on MDH1-CIT1 complex assembly and oxygen consumption rate. (A-D) NanoBiT signal indicating effect of galactose, sucrose, fructose, and glucose on MDH1-CIT1 interaction. Cells were cultured in fresh SD-Raff media in the control condition (black). The cells are treated with 2% sugar application to the SD-Raff-grown cells at 0 min (red). Relative luciferase unit (RLU) was calculated by normalizing the luciferase signals by the average signals during three pre-treatment time points. **(E)** Effects of glucose and inorganic phosphate (fermentation inhibitor) on oxygen consumption rate. Basal O_2_ consumption rate of SD-Raff grown cells was measured. Glucose and inorganic phosphate were added and O_2_ consumption rate was measured for 5 minutes. All data in A-E are presented as mean ± s.d. Statistical differences against the control samples were assessed by Student’s *t*-test at each time point (N=4). Asterisks indicate significant differences with *p*<0.05.

**Figure 2 – figure supplement 3.**
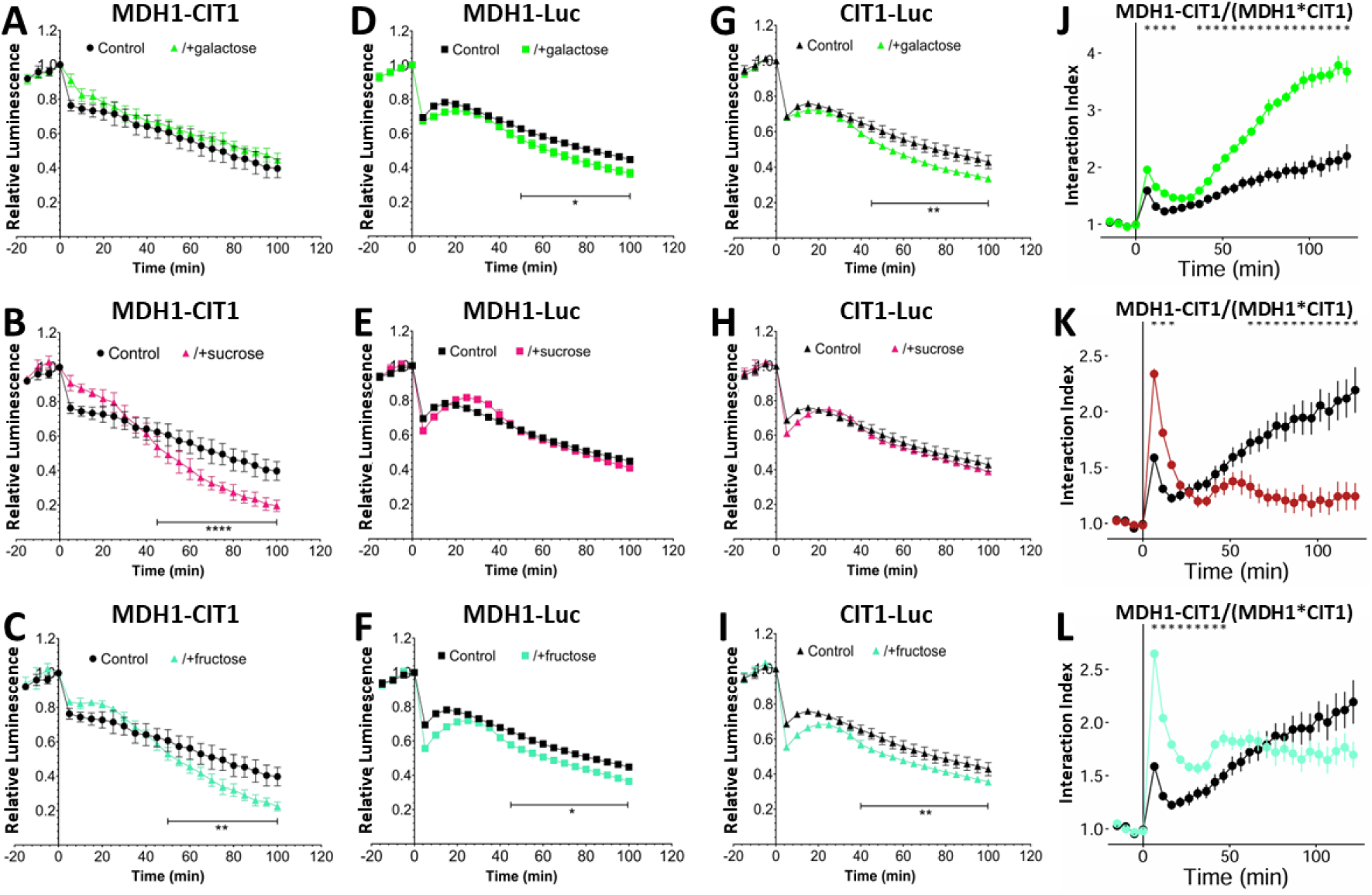
Repeated experiments investigating the effects of sugars on MDH1-CIT1 complex assembly and MDH1 and CIT1 protein abundance following the addition of galactose (green), sucrose (magenta), and fructose (cyan). (A-C) NanoBIT signal indicating effect of galactose **(A)**, sucrose **(B)**, and fructose **(C)** on MDH1-CIT1 interaction. Cells were cultured in fresh SD-Raff media in the control condition (black). The cells are treated with 2% sugar application to the SD-Raff-grown cells at 0 min. Relative luciferase unit (RLU) was calculated by normalizing the luciferase signals by the average signals during three pre-treatment time points. **(D-F)** MDH1 protein levels monitored by the luminescence of MDH1 fused with full-length NanoLUC luciferase. **(G-I)** CIT1 protein levels monitored by the luminescence of CIT1 fused with full-length nanoLUC luciferase. Relative luminescence was calculated by normalizing the luciferase signals by the average signals during three pre-treatment time points. Statistical differences against the control samples were assessed by Student’s *t*-test at each time point (N=4). Asterisks indicate significant differences (*, *p*<0.05 ; **, *p*<0.01 ; ****, *p*<0.0001). **(J-L)** Interaction index calculated by normalizing NanoBiT signal by those of MDH1-NanoLUC and CIT1-NanoLUC.

**Figure 2 – figure supplement 4.**
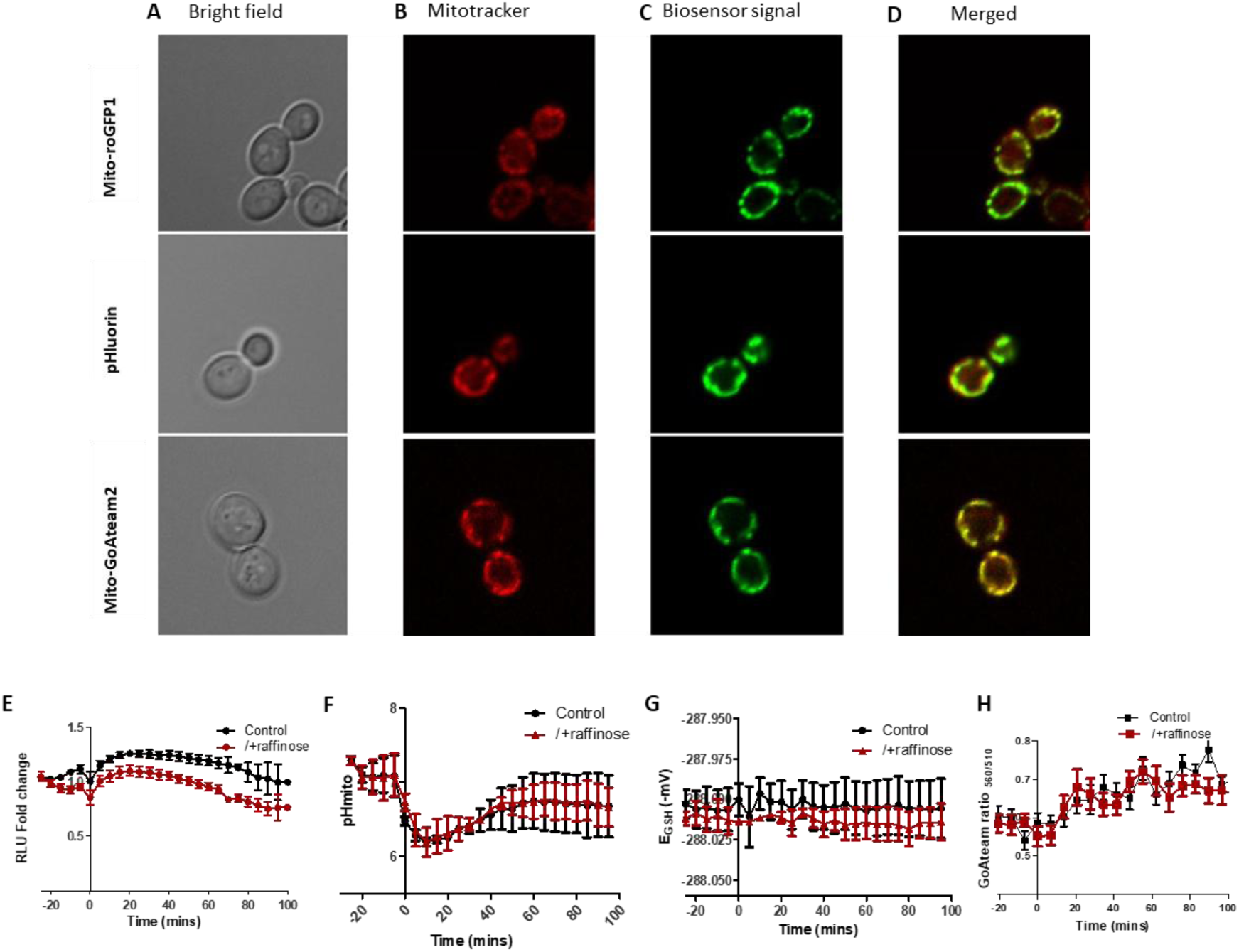
Biosensors indicate mitochondria microenvironments. (A-D) Subcellular localizations of fluorescent biosensors. The yeast strains expressing Mito-roGFP1 (upper panels), pHluorin (middle panels), and Mito-GoAteam2 (lower panels) were observed by fluorescent microscopy in the SD-Raff media. The cells were stained with Mitotracker orange prior to the analysis. **(A)** Bright field image. **(B)** Mito tracker signal. **(C)** Biosensor signals. **(D)** Merged images of the A to C. **(E)** Effect of raffinose addition on MDH1-CIT1 interaction. Cells were cultured in SD-Raff media in the control condition (black). The cells are applied with 2% raffinose to the SD-Raff-grown cells at 0 min (red). Relative luciferase unit (RLU) was calculated by normalizing the luciferase signals by the average signals during three pre-treatment time points. **(F)** Mitochondrial matrix pH in control cells (black) and cells applied with additional raffinose (red). **(G)** Mitochondrial matrix redox state reported as redox potential of roGFP1 (mV). **(H)** Mitochondrial matrix ATP level indicated by the ratio between 560 and 510 nm emission signals of mito-GoATeam2 sensor. Data in E-H are presented as mean ± s.d. (N=4). No statistical difference was detected between the control and raffinose cells.

**Figure 3 – figure supplement 1.**
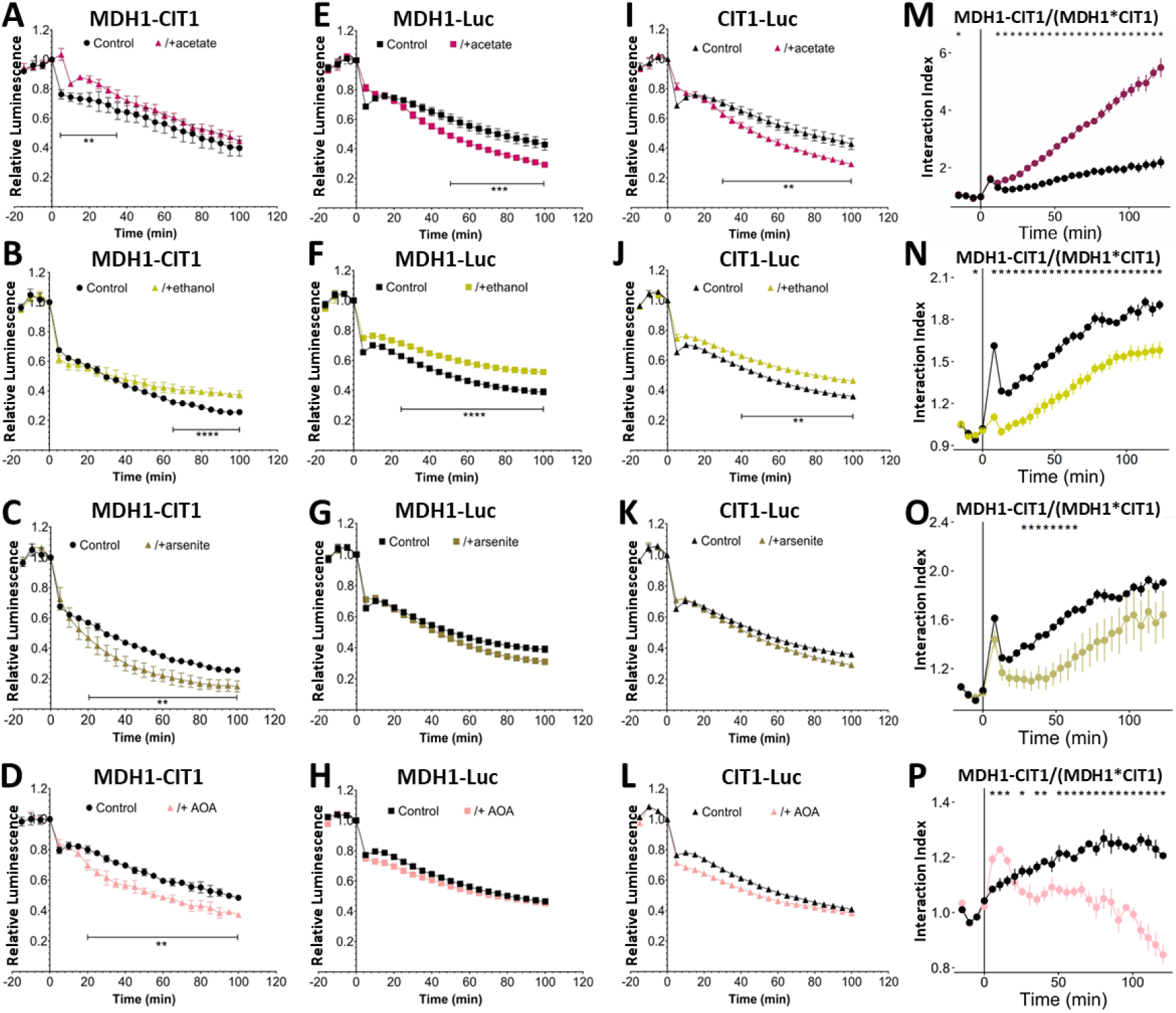
MDH1-CIT1 complex association and MDH1 and CIT1 protein levels following TCA cycle activation and inhibition (repeated experiment). Cells were cultured in SD-Raff media in the control condition (black). The TCA cycle activator (acetate, dark red; ethanol, light yellow) and inhibitors (arsenite, dark yellow; aminooxyacetate, AOA, pink) were applied at 0 min. **(A-D)** NanoBIT signal indicating the MDH1-CIT1 interaction. **(E-H)** MDH1 protein levels monitored by the luminescence of MDH1 fused with full-length NanoLUC luciferase. **(I-L)** CIT1 protein levels monitored by the luminescence of CIT1 fused with full-length NanoLUC luciferase. Relative luminescence Unit (RLU) was calculated by normalizing the luciferase signals by the average signals during three pre-treatment time points. Statistical differences against the control samples were assessed by Student’s *t*-test at each time point (N=4). Asterisks indicate significant differences (**, *p*<0.01; ***, *p*<0.001). **(M-P)** Interaction index calculated by normalizing NanoBiT signal by those of MDH1-NanoLUC and CIT1-NanoLUC. Asterisks indicate time points with statistically significant differences (*p*<0.05) between control and treatment conditions (N=4).

**Figure 4 - figure supplement 1.**
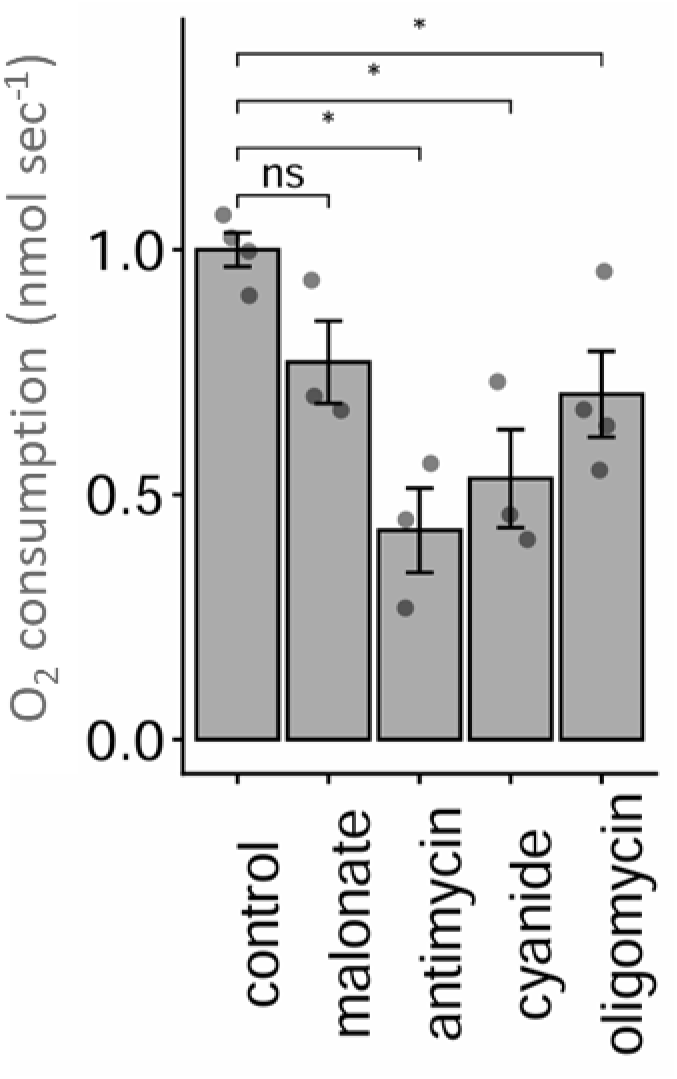
Effect of ETC inhibitors on O_2_ consumption rate. Basal O_2_ consumption rate was measured, then inhibitor was added and O_2_ consumption rate was measured for 5 minutes. Data is presented as mean ± s.d. (N=4), Statistical differences against the control samples were assessed by Student’s *t*-test. Asterisks indicate significant differences with *p*<0.05.

**Figure 4 - figure supplement 2.**
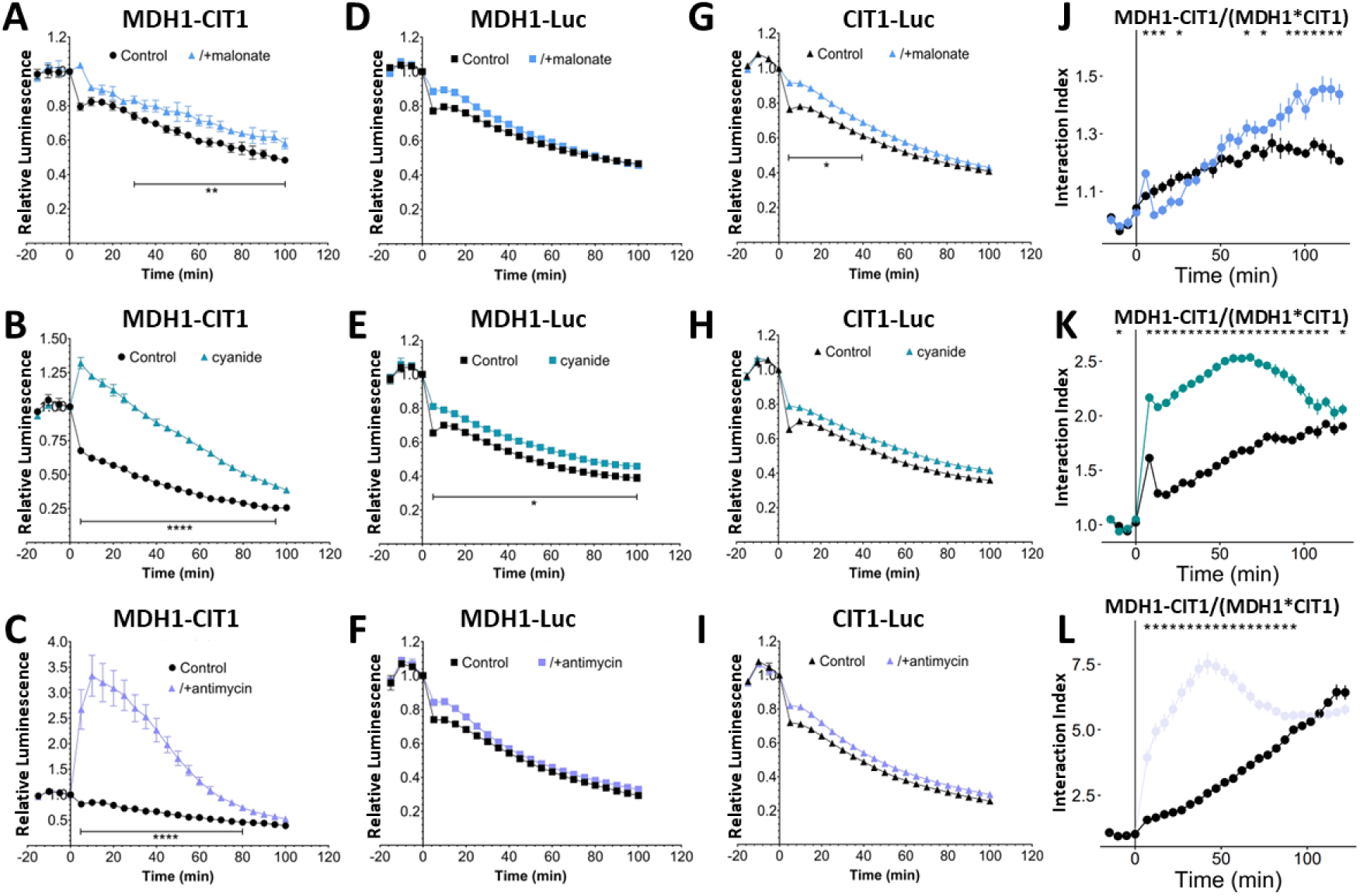
Effect of ETC inhibitors on MDH1-CIT1 complex association and MDH1 and CIT1 protein levels (repeated experiment). Cells were cultured in SD-Raff media in the control condition (black). The electron transport chain inhibitors (malonate, light blue; cyanide, cyan; antimycin, light purple) were applied at 0 min. **(A-C)** NanoBIT signal indicating MDH1-CIT1 interaction. **(D-F)** MDH1 protein levels monitored by the luminescence of MDH1 fused with full-length NanoLUC luciferase. **(G-I)** CIT1 protein levels monitored by the luminescence of CIT1 fused with full-length NanoLUC luciferase. Relative luminescence Unit (RLU) was calculated by normalizing the luciferase signals by the average signals during three pre-treatment time points. Statistical differences against the control samples were assessed by Student’s *t*-test at each time point (N=4). Asterisks indicate significant differences (*, *p*<0.05; **, *p*<0.01; ****, *p*<0.0001). **(J-L)** Relative luminescence of the NanoBiT signal normalized by those of MDH1-NanoLUC and CIT1-NanoLUC. Asterisks indicate the time points with statistical significance (*p*<0.05) between control and treatment conditions.

**Figure 4 – figure supplement 3.**
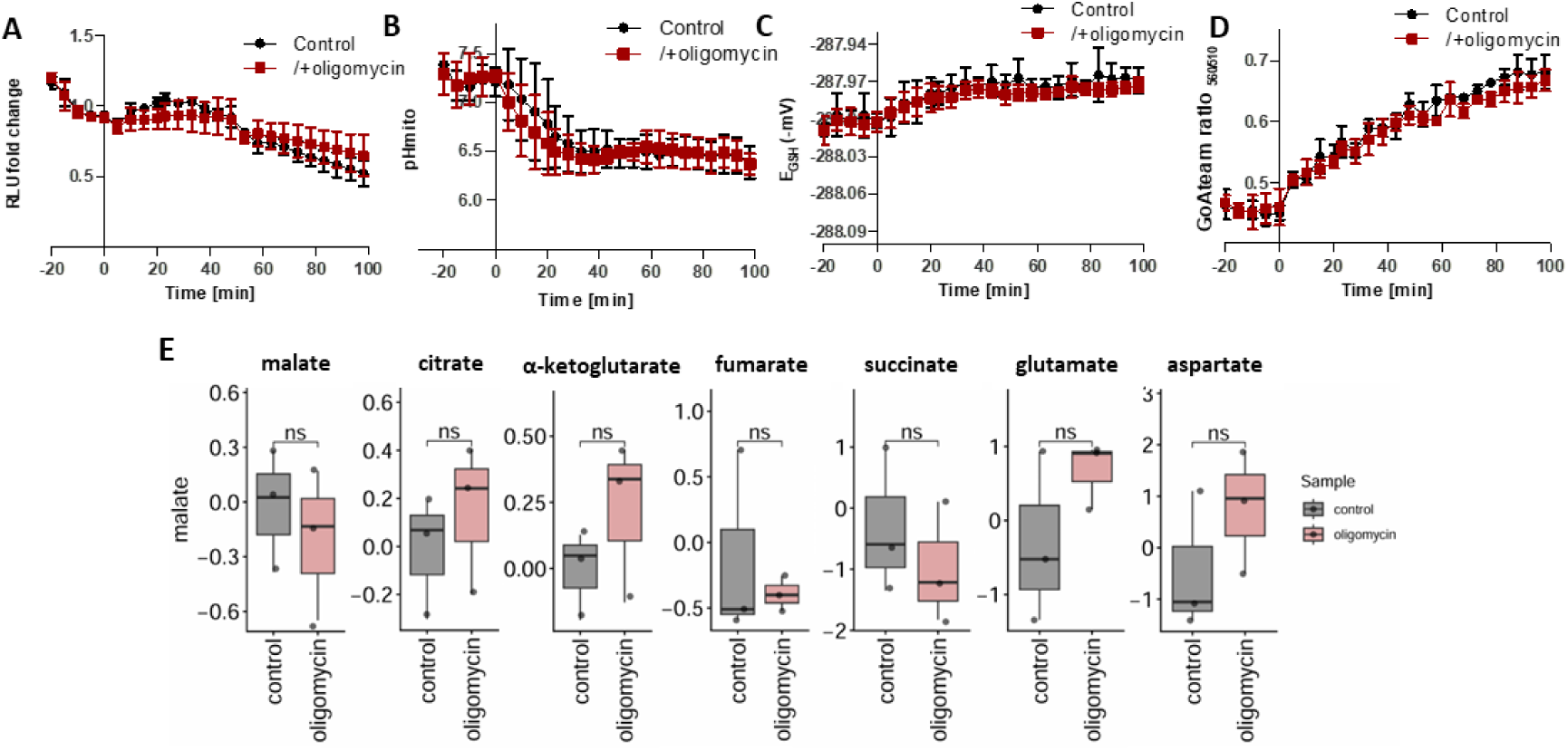
Effects of Complex V inhibition on MDH1-CIT1 complex association, mitochondrial microenvironments, and cellular metabolite levels. Cells were cultured in SD-Raff media in the control condition (black). Oligomycin was applied at 0 min (red). **(A)** NanoBiT signal indicating MDH1-CIT1 interaction. Relative luciferase unit (RLU) was calculated by normalizing the luciferase signals by the average signals during three pre-treatment time points. **(B)** Mitochondrial matrix pH. **(C)** Mitochondrial matrix redox states as GSH/GSSG equivalent (mV). **(D)** Mitochondrial matrix ATP level indicated by the ratio between 560 and 510 nm emission signals of mito-GoATeam2 sensor. All data in A-D are presented as mean ± s.d. (N=4). **(E)** Cellular metabolite levels after 80 min of oligomycin treatment. The boxes, lines, error bars, and points indicate interquartile range, median, minimum, and maximum values, and outliers, respectively. Statistical differences against the control samples were assessed by Student’s *t*-test (N=3). No statistical difference was detected between control and oligomycin-treated cells.

**Figure 4 – figure supplement 4.**
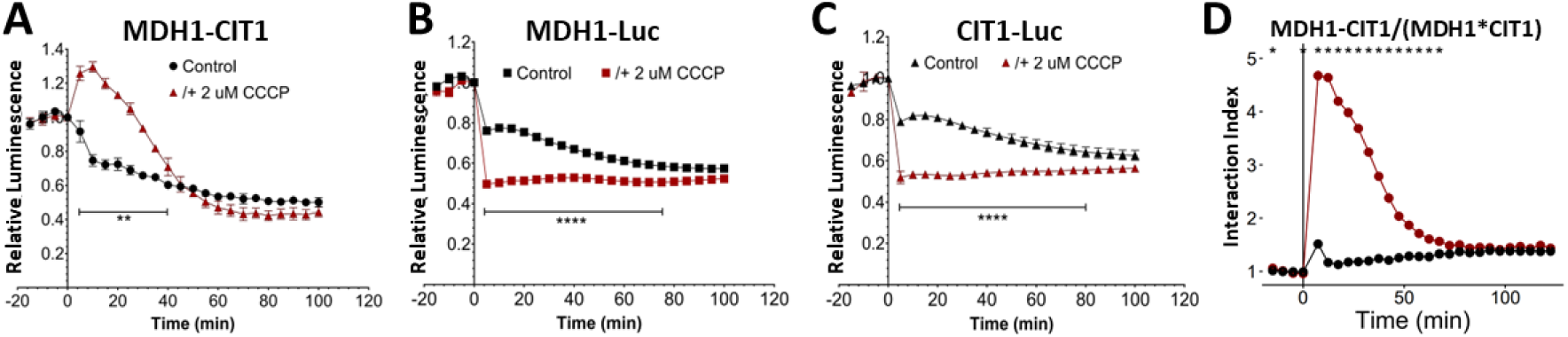
MDH1-CIT1 complex association and MDH1 and CIT1 protein levels following oxidative phosphorylation uncoupler (carbonyl cyanide 3-chlorophenylhydrazone; CCCP) application. Cells were cultured in SD-Raff media in the control condition (black). CCCP (dark red) were applied at 0 min. **(A)** NanoBIT signal indicating MDH1-CIT1 interaction. **(B)** MDH1 protein levels monitored by the luminescence of MDH1 fused with full-length NanoLUC luciferase. **(C)** CIT1 protein levels monitored by the luminescence of CIT1 fused with full-length NanoLUC luciferase. Relative luminescence Unit (RLU) was calculated by normalizing the luciferase signals by the average signals during three pre-treatment time points. Statistical differences against the control samples were assessed by Student’s *t*-test at each time point (N=4). Asterisks indicate significant differences (**, *p*<0.01 ; ****, *p*<0.0001). **(D)** Interaction index calculated by normalizing NanoBiT signal by those of MDH1-NanoLUC and CIT1-NanoLUC. Asterisks indicate time points with statistically significant differences (*p*<0.05) between control and CCCP conditions (N=4).

**Figure 5 – figure supplement 1.**
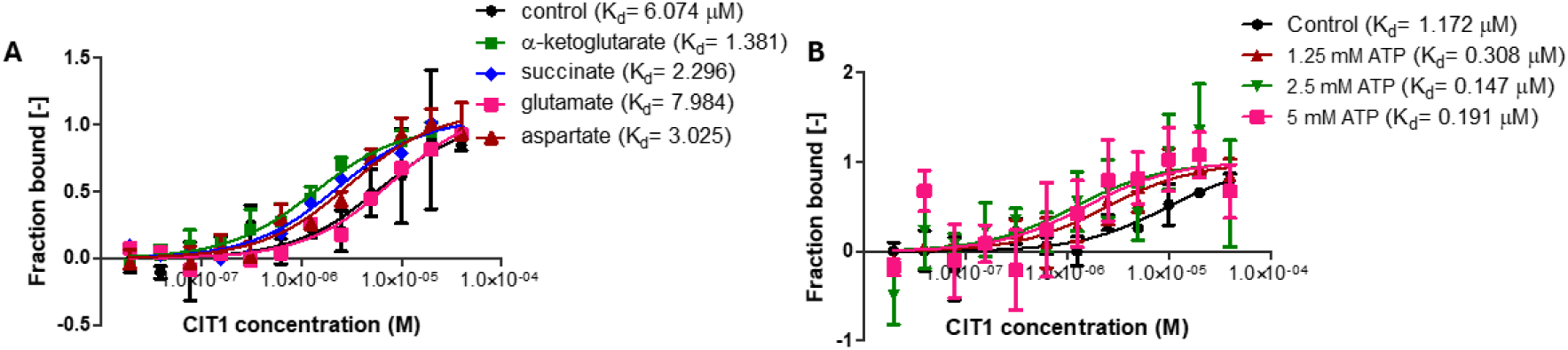
**e**The affinity of the MDH1-CIT1 multienzyme complex was analyzed by microscale thermophoresis (MST) using fluorescently labeled MDH1 as the target and CIT1 as the ligand. Curves represent the response (fraction bound) against CIT1 concentration. Points represent the means of fraction bound, and the error bars represent the standard deviations of three measurements (N=3). **(A)** Effects of metabolites. The MDH1-CIT1 interaction was determined in the buffer (control; black) with 10 mM α-ketoglutarate (green), 10 mM succinate (blue), 10 mM glutamate (pink), and 10 mM aspartate (dark red). **(B)** Effects of 1.25 mM (brown), 2.5 mM (green), and 5 mM (pink) ATP. The *K*_d_ values of MDH1-CIT1 interaction were shown next to the legend.

## Supplementary Dataset 1 (separate file)

**Metabolite profiling data of the *S*. *cerevisiae* cells.** The relative metabolite levels were used for the analyses in this study. The raw peak heights, quantitative ion m/z, and the retention time of the each analyzed peak are also indicated. The metabolite profiling was conducted in two batchs. CONTROL1, GLUCOSE, CYANIDE, ACETATE, ARSENITE, and AOA conditions were analyzed in the first batch (N=7). CONTROL2, MALONATE, OLIGOMYCIN, and ANTIMYCIN were analyzed in the second batch (N=3).

