## Supplementary Method for "Dynamic assembly of malate dehydrogenase-citrate synthase multienzyme complex in the mitochondria"

**General procedures**

PCR reactions for the construction of vectors and integration cassettes were conducted with Phusion High-Fidelity DNA polymerase (Thermo Fisher Scientific, Waltham, MS, USA). DreamTaq Green DNA polymerase (Thermo Fisher Scientific) was used for the amplification of vector cloning sites and yeast genome to verify the integration of desired sequences. The Sanger sequencing of the PCR products was conducted by Eurofins Scientific (Luxembourg City, Luxembourg). Restriction digestion and ligation were conducted by manufacturer's protocols using the enzymes from New England Biolab (NEB, Ipswich, MA, USA). Chemically competent *E. coli* DH5α was used for vector construction. Frozen-EZ Yeast Transformation Kit (Zymo Research Corporation, CA, USA) was used for yeast cell transformation. All oligo DNAs were synthesized by MilliporeSigma (Burlington, MA, USA) and Eurofins Scientific. Sequences of the primers used in this study are in Supplementary Table 1.

**Generation of the nanoBiT and nanoLUC reporter yeast strains**

**Vector construction**

The DNA fragments for genomic DNA recombination were constructed in the pGDBα1 vector using the Golden Braid technology (Sarrion-Perdigones et al., 2013). The procedure is summarized in Figure. First, the cloning site in the pDGB2Ω2 vector (Sarrion-Perdigones et al., 2013) was replaced to generate three vectors containing desired 4-base overhang by BsaI digestion. The pDGB2Ω2 cloning site was amplified using the primers with modified overhang sequences and EcoRV recognition sequences on both ends (pGDB2Ω2-1, pGDB2Ω2-2, and pGDB2Ω2-3 primer sets; Supplementary Table 1). Purified PCR products were cloned into pGDB2Ω2 at the EcoRV site by T4 ligase, resulting in the pGDB2Ω2-1, pGDB2Ω2-2, and pGDB2Ω2-3 vectors including GGAG/TCCT, TCCT/CGTC, CGTC/CGCT BsaI restriction cites, respectively (Figure). The inserted sequences were amplified using pDGB1seq primer sets and the sequences were verified.

Second, the vectors harboring 5' half of tag-coding sequence, URA3 selection marker, and 3' half of tag-coding sequence were generated. The DNA sequence coding linker-HA-nanoBiT large subunit (LgBiT-HA), linker-cMyc-nanoBiT small subunit (SmBiT-cMyc), and linker-HA-full length nanoLUC luciferase (nLUC-HA) were synthesized by Genewiz (Azenta Life Sciences, South Plainfield, NJ, USA). The sequences of the synthesized DNA are in Supplementary Table 2. The 5' and 3' halves of tag-coding sequences were amplified using the primers with 5' extensions containing BsaI recognition sequences to generate GGAG/TCCT and CGTC/CGCT overhangs to be cloned into the BsaI sites of pGDB2Ω2-1 and pGDB2Ω2-3, resulting in pGDB2Ω2-5' and pGDB2Ω2-3' vectors, respectively. pDGB2Ω2-LgBiT5'and pDGB2Ω2-LgBiT3' primer sets were used to amplify the 5' and 3' halves of the large nanoBiT subunits using LgBit-HA as the template to construct pDGB2Ω2-5'LgBiT and pDGB2Ω2-3'LgBiT vectors, respectively. pDGB2Ω2- SmBiT 5'and pDGB2Ω2- SmBiT 3' primer sets were used to amplify the 5' and 3' halves of the small nanoBiT subunits using SmBit-cMyc as the template to construct pDGB2Ω2-5'SmBiT and pDGB2Ω2-3'SmBiT vectors, respectively. Full length nanoLUC luciferase sequence was amplified using pDGB2Ω2-nLUC5' and pDGB2Ω2-nLUC3' primer sets with nLUC-HA as the template to construct pDGB2Ω2-5'nLUC and pDGB2Ω2-3'nLUC vectors, respectively. URA3 gene was amplified from genomic DNA of *Kluyveromyces lactis* using the primers with BsaI sites with TCCT/CGTC sequences (pDGB2Ω2-Ura primer set) and cloned into pGDB2Ω2-2 to construct the pGDB2Ω2-URA3 vector. The inserted sequences were amplified using the pDGB1seq primer set and their sequences were verified.

Third, the DNA fragments were assembled into the pGDBα1 vector using the Golden Braid procedure. The full length nanoLUC, LgBiT and SmBiT coding sequences were integrated at direct downstream of the Mdh1 (chrIV:3300230) and Cit1 (chrX:303993) genes on the BY4741 genome for C-terminal fusion. pGDB2Ω2-5', pGDB2Ω2-URA3, pGDB2Ω2-3' (300 ng each), and 200 ng pGDBα1 were incubated with 20U BsaI-HF (NEB) and 400U T4DNA ligase (NEB) at 37°C 4 min, 16°C 5 min, and 50°C 5 min for 28 cycles. The reaction was terminated by heating at 80°C for 5 min after the cycles. The reaction mixtures were used to transform *E. coli* and the colonies were selected on LB plates containing 50 μg/ml kanamycin, 20 μg/ml Xgal (GoldBio) and 25 μg/ml IPTG (GoldBio). The plasmids were extracted from the white colonies. Their inserted sequences were amplified using the pDGB1seq primer set and their sequences were verified by sequencing. pGDB2Ω2-5'LgBiT, pGDB2Ω2-URA3, and pGDB2Ω2-3'LgBiT were used to construct pGDBα1-LgBiT vector. pGDB2Ω2-5'SmBiT, pGDB2Ω2-URA3, and pGDB2Ω2-3'SmBiT vectors were used to construct pGDBα1-SmBiT vector. pGDB2Ω2-5'SmBiT, pGDB2Ω2-URA3, and pGDB2Ω2-3'SmBiT vectors were used to construct pGDBα1-SmBiT vector. pGDB2Ω2-5'nLUC, pGDB2Ω2-URA3, and pGDB2Ω2-3'nLUC vectors were used to construct pGDBα1-nLUC vector.

**Scarless integration of tagging sequences into yeast genome**
The integration cassettes were PCR-amplified using the primers annealing directly downstream of the BsaI sites with 37-40 base 5' extension complementary to the franking sequences of the genomic insertion sites. The fragment to integrate small nanoBiT subunit into the downstream of MDH1 gene was amplified using MDH1-SmBiT primer set with pGDBα1-SmBiT vector as the template. The fragment to integrate large nanoBiT subunit into the downstream of CIT1 gene was amplified using CIT1-LgBiT primer set with pGDBα1-LgBiT vector as the template. The fragment to integrate full length nanoLUC coding sequence into the downstream of MDH1 gene was amplified using MDH1-nLUC primer set with pGDBα1-nLUC vector as the template. Yeast cells were transformed with the PCR products and the strains with integrated sequences were selected on URA- plates. The genomic DNA was isolated from the positive clones and the tag sequence insertion was assessed by PCR using the primers annealing the adjacent region of the insertion sites (MDH1seq and CIT1seq primer sets) and the URA3 coding sequence (Ura3intFw/MDH1seqRv and Ura3intFw/CIT1seqRv primer sets).
The URA3 marker was ejected by endogenous homologous recombination between the repeats in the inserted sequence. The verified transformants were cultured in liquid YPD media containing uracil for two days. The strains with removed URA3 marker were selected on the URA-containing SD plate with 1 mg $mL^{-1}$ 5-fluoroorotic acid (5-FOA). The lack of URA3 maker was assessed by PCR using MDH1seq and CIT1seq primer sets. Correct integration of the inserted fragments was confirmed by sequencing.

**Recombinant MDH1 and CIT1 protein production**
**Vector construction**
The Cit1 and Mdh1 coding sequences without the mitochondrial targeting signal were amplified from *S. cerevisiae* BY4741 cDNA library generated by SuperScript III First-Strand Synthesis System (Invitrogen, Thermo Fisher Scientific) using the cit1-Fw/cit1-Rv and mdh1-Fw/mdh1-Rv primer sets (Supplementary Table 1). The cDNA fragments were cloned into the NdeI/XhoI restriction site in the expression plasmid pET21b with a c-terminal 6x His tag, using the ClonExpress II One Step Cloning kit (Nanjing Vazyme Biotech Co. Ltd., China) following the manufacturer's instructions. The resulting plasmids were extracted, and the inserts were amplified by cit1-verFw/ctmd-verRv and mdh1-verFw/ctmd-verRv primer sets (Supplementary Table 1) to verify the inserted sequences. The plasmids for the expression of CIT1 (pET-cit1) and MDH1 (pET-mdh1) were transformed into chemically competent *E. coli* BL21 (DE3) cells.

**Protein expression and purification**
The *E. coli* BL21 (DE3) strains harboring pET-cit1 and pET-mdh1 were cultivated in terrific broth (TB) medium at 37°C with shaking at 220 rpm to $OD_{600}$= 0.6–0.8, and 0.8 and 0.2 mM of isopropyl-β-D-thiogalactopyranoside (IPTG) was applied for CIT1 and MDH1 expression induction, respectively. The cultures were incubated for an additional 20 h at 18 °C, and the cells were harvested by centrifugation at 6000 x *g* and 4 °C for 20 min. The cell pellets were resuspended in cell lysis buffer (50 mM tris-HCl buffer, pH 8.0, 500 mM NaCl, 10% glycerol, 20 mM imidazole, 1 tablet of cocktail of EDTA-free protease inhibitors/100 ml, 200 μg/ml lysozyme, 20 μg/ml DNase, 1 mM $MgCl_2$, 0.5% Tween 20 and 10 mM DTT) using 5-fold volume of the weight of cell pellets. The cells were then disrupted by sonication using Sonic Dismembrator 550 (Fisher Scientific, Hampton, NH, USA) with the amplitude set at 6, pulser on 5s, pulser off 15s and a total process time of 7 min. Following the centrifugation at 16,000 x *g* at 4 °C for 70 min, buffer), recombinant proteins in the supernatants were purified using Ni-NTA His SpinTrap

columns (GE Healthcare, Chicago, IL, USA), and the proteins were eluted with concentration-gradient imidazole in protein purification buffer (50 mM tris-HCl buffer, pH 8.0, 500 mM NaCl, 10 mM β-mercaptoethanol, 20 mM imidazole (binding buffer), 40 mM, 60 mM and 80 mM imidazole (wash buffers), and 500 mM imidazole (elution buffer)). The purified enzymes were desalted using an Amicon Ultra-4 centrifugal concentrator (10 kDa MWCO, MilliporeSigma) at 4 °C to exchange to buffer A (50 mM tris-HCl buffer, pH 8.0). Enzyme purity was estimated using SDS–PAGE with coomassie blue staining. Glycerol was added to the purified proteins to a final concentration of 10%. The purified proteins were then flash-frozen with liquid nitrogen and stored at -80 °C until further use. Protein concentration was determined using Bio-Rad Protein Assay Kit (Bio-Rad Laboratories, Hercules, CA, USA) following the manufacturer's instructions. The enzyme activity was verified as described in the Method section.

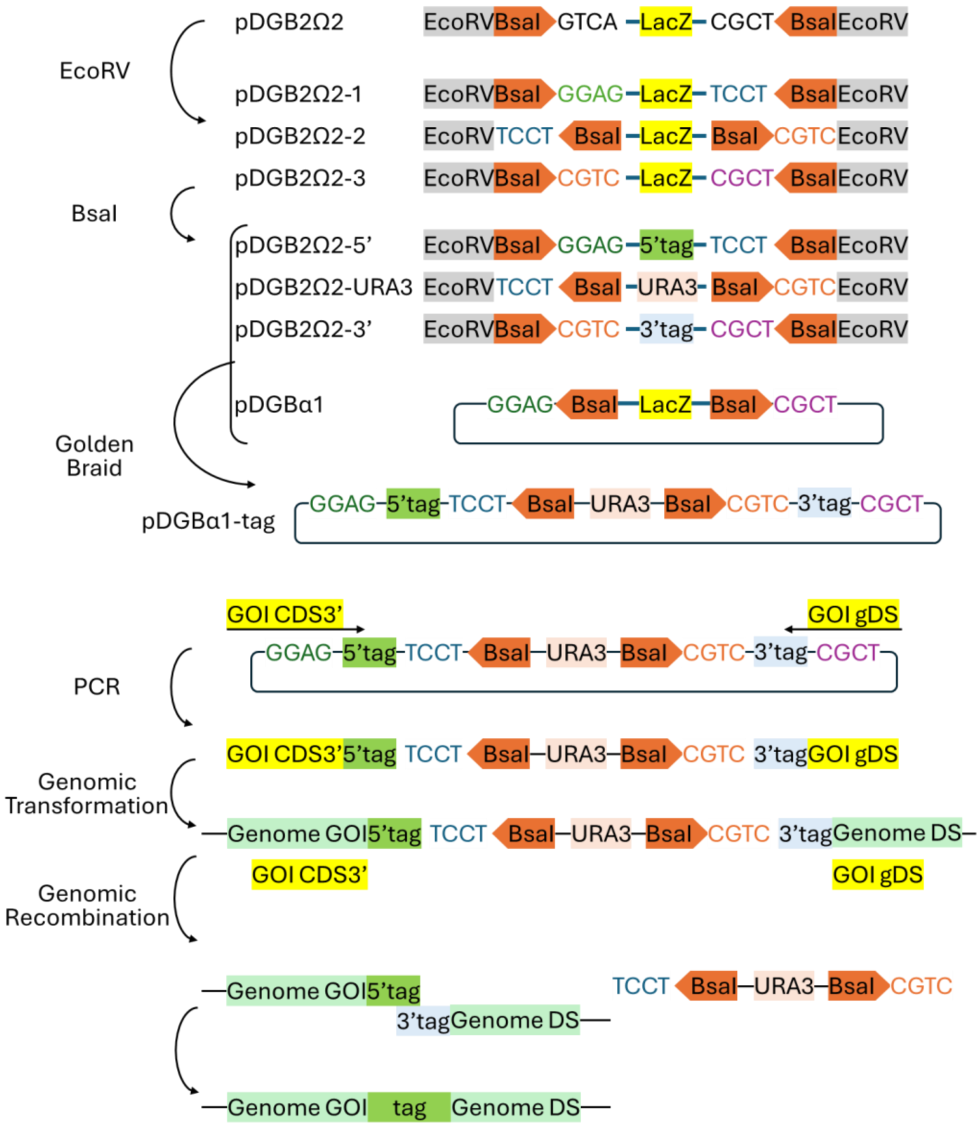

**Figure. Schematics of vector construction and genomic integration procedures.** Colored texts indicate the overhang sequences of the BsaI digestion sites. The “tag” indicates the integrated tag-coding sequences, namely large nanoBiT subunit, small nanaBiT subunit, and nanoLUC luciferase. GOI, gene of interest; CDS, coding sequence; gDS, genomic downstream sequence; Genome, genomic sequence.

**Supplementary Table 1. Primers used in this study.** Fw and Rv indicate forward and reverse primers, respectively. Bold letters indicate the 4-base overhang in the BsaI digestion sites. Underlined sequences indicate plasmid-annealing sequences.

| Table S1 | Primers used in this study |
|---|---|
| Primer Name | Primer Sequence |
| pGDB2Ω2-1Fw | TCATGATATCGGTCTCA**GGAG**CACAGCTTGTCTGTAAGCG |
| pGDB2Ω2-1Rv | CATTGATATCGGTCTCA**AGGA**CAGCTGGCACGACAGGTTTC |
| pGDB2Ω2-2Fw | GATATC**TCCT**AGAGACCCACAGCTTGTCTGTAAGCGG |
| pGDB2Ω2-2Rv | CATTGATATC**GACG**AGAGACCCAGCTGGCACGACAGGTTTC |
| pGDB2Ω2-3Fw | CATTGATATCGGTCTCA**CGTC**CACAGCTTGTCTGTAAGCGG |
| pGDB2Ω2-3Rv | CATTGATATCGGTCTCA**AGCG**CAGCTGGCACGACAGGTTTC |
| pDGB2Ω2-LgBiT5’Fw | CATTCATTGGTCTCA**GGAG**CAGCTTGTAAGATCCCAAACG |
| pDGB2Ω2-LgBiT5’Rv | GTTGTATTGGGTCTCA**AGGA**GTAACACCATCAATAACTAAAGTACCA |
| pDGB2Ω2-UraFw | GGTGTTAC**TCCT**TGAGACCCAATACAACAGATCACGTG |
| pDGB2Ω2-UraRv | AGCAGTTT**GACG**TGAGACCCGTTTTATTTAGGTTCTATCGAGG |
| pDGB2Ω2-LgBiT3’Fw | CATTACGGTCTCA**CGTC**CAAACTGCTGCT |
| pDGB2Ω2-LgBiT3’Rv | CTAGAGTAGGTCTCT**AGCG**GCTAAAATACGTTCACATAAACGCCAACC |
| pDGB2Ω2-SmBiT5’Fw | ATAGGTCTCA**GGAG**GCATGTAAAATTCCTAATGATTTAA |
| pDGB2Ω2-SmBiT5’Rv | GAACTAGGTCTCA**AGGA**TAAAATTTCTTCAAATAAACGATAACCAG |
| pDGB2Ω2-SmBiT3’Fw | CATTACGGTCTCA**CGTC**GCATGTAAAATTCC |
| pDGB2Ω2-SmBiT3’Rv | CATAAGGGTCTCT**AGCG**TTATAAAATTTCTTCAAATAAACGATAA |
| pDGB2Ω2-nLUC5’Fw | CATTCATTGGTCTCA**GGAG**CAGCTTGTAAGATCCCAAACG |
| pDGB2Ω2-nLUC5’Rv | GTTGTATTGGGTCTCA**AGGA**GTAACACCATCAATAACTAAAGTACCA |
| pDGB2Ω2-nLUC3’Fw | ATAAAACGGGTCTCA**CGTC**AAACTGCTGGTTATAATTTAGATCAAG |
| pDGB2Ω2-nLUC3’Rv | CTAGAGTAGGTCTCT**AGCG**GCTAAAATACGTTCACATAAACGCCAACC |
| MDH1-SmBiT-Fw | AAGAATATCGAAAAGGGTGTCAACTTTGTTGCTAGTAAAGCATGTAAAATT CCTAATGA |
| MDH1-SmBiT-Rv | TTTTTTTCCCTATTTTTCACTCTATTTCTGATCTTATAAAATTTCTTCAAA TAAACG |
| CIT1-LgBiT-Fw | AATACAAGGAGTTGGTAAAGAAAATCGAAAGTAAGAACGCTTGTAAGATCC CAAACGAC |
| CIT1-LgBiT-Rv | TGAAAATACGTGTTTGAATAGTCGCATACCCTGAATCTCGTGTTACTATTA ATTCTTA |
| MDH1-nLUC-Fw | AAGAATATCGAAAAGGGTGTCAACTTTGTTGCTAGTAAAAGACCAGCTTGT AAGATCCCA |
| MDH1-nLUC-Rv | TTCCCTATTTTTCACTCTATTTCTGATCTTGAACAATTTAAGCTAAAATAC GTTCACA |
| MDH1seqFw | GCATCTCCGGTCACTTTGGG |
| MDH1seqRv | CTAGTTGATTTTTGGCAGTTTCCTTCCTTTC |
| CIT1seqFw | GCCAGAGCTATTGGTGTGTTACC |
| CIT1seqRv | CGGTAGGCATAGGGGACTCAAAG |
| Ura3intFw | GCGTTACCACCATCCAATGCAGAC |
| pDGB1seqFw | CGCCAGCAACGCGGCCTT |
| pDGB1seqRv | GCAAGCGGTTGCCCACCGTC |
| cit1-Fw | <u>CTTTAAGAAGGAGATATACATATG</u>AGTAGCGCCTCCGAACAAACG |
| cit1-Rv | <u>AGTGGTGGTGGTGGTGGTGCTCGAG</u>GTTCTTACTTTCGATTTTCTTTACCA ACTCCTTG |
| mdh1-Fw | <u>CTTTAAGAAGGAGATATACATATG</u>TATAAAGTGACTGTTTTGGGTGCAGGC |
| mdh1-Rv | <u>AGTGGTGGTGGTGGTGGTGCTCGAG</u>TTTACTAGCAACAAAGTTGACACCCT TTTC |
| cit1-verFw | GGGCTACGAAAACAAGGATTTTATTGAC |
| mdh1-verFw | CATCAACGCAAGCATCGTTC |
| ctmd-verRv | GTTATGCTAGTTATTGCTCAGCGGTG |

**Supplementary Table 2. Sequences of the synthesized DNA.**

| Table S2 | Sequences of the synthesized DNA |
|---|---|
| DNA name | Sequences |
| nLUC-HA | CCCGGGAGACCAGCTTGTAAGATCCCAAACGACTTGAAGCAAAAGGTTA<br>TGAACCACTACCCATACGACGTACCAGATTACGCTATGGTTTTTACTTT<br>AGAAGATTTTGTTGGTGATTGGCGTCAAACTGCTGGTTATAATTTAGAT<br>CAAGTTTTAGAACAAGGTGGTGTTTCTTCTTTATTTCAAAATTTAGGTG<br>TTTCTGTTACTCCTATTCAACGTATTGTTTTATCTGGTGAAAATGGTTT<br>AAAAATTGATATTCATGTTATTATTCCTTATGAAGGTTTATCTGGTGAT<br>CAAATGGGTCAAATTGAAAAAATTTTTAAAGTTGTTTATCCTGTTGATG<br>ATCATCATTTTAAAGTTATTTTACATTATGGTACTTTAGTTATTGATGG<br>TGTTACTCCTAATATGATTGATTATTTTGGTCGTCCTTATGAAGGTATT<br>GCTGTTTTTGATGGTAAAAAAATTACTGTTACTGGTACTTTATGGAATG<br>GTAATAAAATTATTGATGAACGTTTAATTAATCCTGATGGTTCTTTATT<br>ATTTCGTGTTACTATTAATGGTGTTACTGGTTGGCGTTTATGTGAACGT<br>ATTTTAGCTGTCGAC |
| LgBiT-HA | GGATCCAGACCAGCTTGTAAGATCCCAAACGACTTGAAGCAAAAGGTTA<br>TGAACCACTACCCATACGACGTACCAGATTACGCTATGGTTTTTACTTT<br>AGAAGATTTTGTTGGTGATTGGGAACAAACTGCTGCTTATAATTTAGAT<br>CAAGTTTTAGAACAAGGTGGTGTTTCTTCTTTATTGCAAAATTTAGCTG<br>TTTCTGTTACTCCTATTCAACGTATTGTTAGATCTGGTGAAAATGCTTT<br>AAAAATTGATATTCATGTTATTATTCCTTATGAAGGTTTATCTGCTGAT<br>CAAATGGCTCAAATTGAAGAAGTTTTTAAAGTTGTTTATCCTGTTGATG<br>ATCATCATTTTAAAGTTATTTTACCATATGGTACTTTAGTTATTGATGG<br>TGTTACTCCTAATATGTTGAATTATTTTGGTCGTCCTTATGAAGGTATT<br>GCTGTTTTTGATGGTAAAAAAATTACTGTTACTGGTACTTTATGGAATG<br>GTAATAAAATTATTGATGAACGTTTAATTACACCTGATGGTTCTATGTT<br>ATTTCGTGTTACTATTAATTCTTAAGATATC |
| SmBiT-cMyc | GATATCAGACCAGCATGTAAAATTCCTAATGATTTAAAACAAAAAGTTA<br>TGAATCATATGGAACAAAAGTTGATTTCTGAAGAAGATTTGGTTACTGG<br>TTATCGTTTATTTGAAGAAATTTTATAAAAGCTT |
